# TGFβ-induced SMAD1/5 activation drives transient epithelial-to-mesenchymal transition in trophoblasts

**DOI:** 10.1101/2025.07.21.665880

**Authors:** Sophia Mähr, Hallfridur Ingolfsdottir, Marta Sorokina Alexdottir, Sunhild Hartmann, Olivia Nonn, Johanna Gunnarsdottir, Þóra Steingrimsdottir, Jerome Jatzlau, Petra Knaus, Gudrun Valdimarsdottir

## Abstract

Successful human placental development requires fetal trophoblasts to undergo epithelial-to-mesenchymal and mesenchymal-to-endothelial transitions, enabling invasion and remodeling of maternal spiral arteries for proper placental perfusion. These trans-differentiation processes are disrupted in preeclampsia, a major cause of maternal and fetal morbidity worldwide. Despite its clinical significance, the molecular mechanisms governing these differentiation trajectories at the fetal-maternal interface remain poorly defined. While Transforming Growth Factor-beta (TGFβ) has been implicated in trophoblast invasion, its role remains controversial. In this study, we report increased SMAD2/3 phosphorylation (pSMAD2/3), but reduced pSMAD1/5 and SNAIL levels in preeclamptic placentas. Mechanistically, we show that TGFβ1 induces transient SMAD1/5 phosphorylation via dual engagement of the TGFβ type I receptor ALK5 and the BMP type I receptor ALK2 in both HTR8/SVneo cells and human trophoblast stem cells. Moreover, single-nucleus transcriptomics of first-trimester placentas revealed co-expression of *ACVR1*, *TGFBR1*, *SMAD1* and *SMAD5* at the transition zone between cytotrophoblasts and extravillous trophoblasts. Our findings suggest that transient TGFβ-induced ALK5/ALK2-SMAD1/5 signaling is critical for EMT initiation, a prerequisite for proper placental differentiation.

## INTRODUCTION

A key process in placental development is the remodelling of uterine spiral arteries as they convert from high-resistance to low-resistance blood vessels to ensure adequate perfusion of the placenta. Between weeks 6 and 12 of gestation, cytotrophoblasts derived from the trophectoderm differentiate into extravillous trophoblasts (EVTs), which invade the maternal decidua and extracellular matrix (ECM) towards the spiral arteries [1–4]. Upon reaching these vessels, EVTs displace vascular smooth muscle cells (vSMC) and replace the maternal endothelium in a process termed pseudovasculogenesis [4, 5]. These invasive and remodelling behaviors require sequential epithelial-to-mesenchymal transition (EMT) and mesenchymal-to-endothelial transition (MEndT), respectively. In preeclampsia, a leading cause of maternal and fetal morbidity, these key processes are disrupted. Shallow trophoblast invasion and incomplete spiral artery remodelling are hallmarks of severe preeclampsia, particularly in early-onset cases (<34 weekśgestation), that can cause placental hypoperfusion, maternal hypertension and proteinuria (**Fig.1A**) [6, 7]. While dysregulated angiogenic signalling has been implicated, the precise molecular mechanisms driving trophoblast differentiation and invasion remain incompletely understood [4, 8, 9].

**Figure 1:**
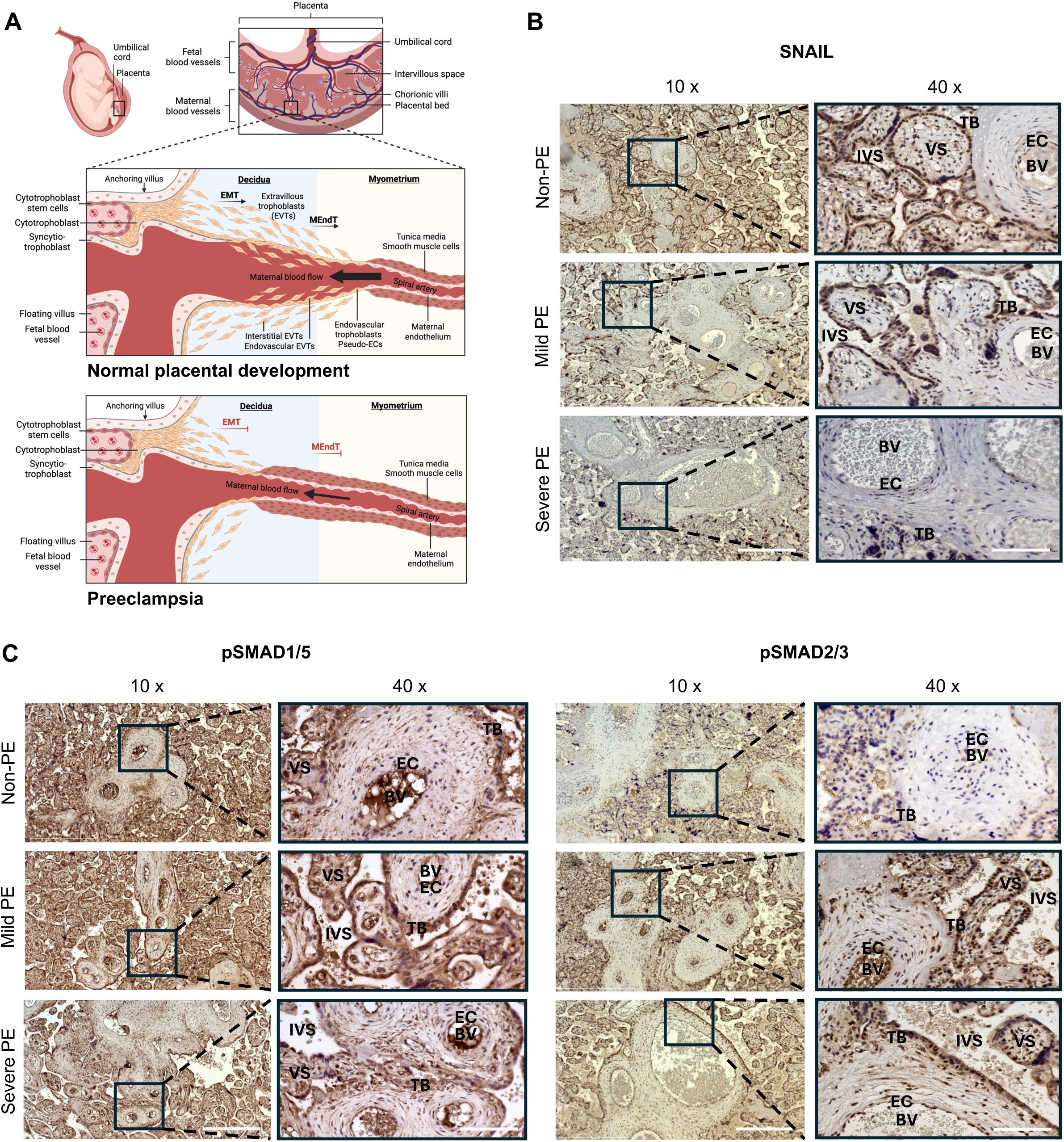
SMAD1/5 phosphorylation and SNAIL expression are reduced in preeclamptic placental samples. **A,** Cytotrophoblasts (CTBs) undergo epithelial-to-mesenchymal transition (EMT) to differentiate into extravillous trophoblasts (EVTs) and invade the extracellular matrix towards the maternal blood vessels. In turn, trophoblasts replace the endothelium and remove vascular smooth muscle cells (vSMCs), a process called pseudovasculogenesis which is caused by mesenchymal-to-endothelial transition (MEndT). In consequence, a high-resistance to low-resistance spiral artery remodeling can occur and ensures a sufficient blood flow and thereby an adequate perfusion of the placenta and the fetus. In preeclampsia, the differentiation processes seem to fail, leading to a shallow trophoblast invasion and reduced transformation of uterine spiral arteries which can cause hypertension and proteinuria as defining symptoms of the disease. **B,C**, Immunohistochemical stainings of SNAIL (**B**), pSMAD1/5 and pSMAD2/3 (**C**) (counterstained with haematoxylin) on placental sections from healthy (Non-PE, n = 5) and preeclamptic women (PE) with mild (n = 5) or severe (n = 5) features. Immunostained sections were imaged with a light microscope using 10x and 40x magnification. Scale bar, 500 µm (10x) and 125 µm (40x). BV – blood vessel, EC – endothelial cells, TB – trophoblast cells, VS – villous space, IVS – intervillous space.

EMT is characterized by the loss of cell polarity and adhesion, accompanied by the acquisition of mesenchymal morphology and motility [10, 11]. These changes are orchestrated by transcription factors, such as SNAIL (*SNAI1*), SLUG (*SNAI2*), ZEB and basic helix-loop-helix (bHLH) families, which repress epithelial markers like E-Cadherin (*CDH1*) and induce mesenchymal markers such as N-Cadherin (*CDH2*) and Vimentin (*VIM*) [12–14]. In the placenta, these EMT regulators are essential for EVT lineage commitment and invasive capacity [15].

Trophoblast cells and cancer cells share several characteristics including proliferation, migration, invasion and immune tolerance. Based on their common features, it is conceivable that both cell types undergo similar molecular mechanisms to induce distinct cellular responses such as EMT [16, 17]. One of the most studied inducers of EMT in both contexts is the TGFβ family [12, 18]. TGFβ signaling is crucial for embryonic development and tissue homeostasis, but its dysregulation has been implicated in tumorigenesis and defective placental development [4, 19].

The TGFβ family consists of structurally related polypeptide growth factors including bone morphogenetic proteins (BMPs), growth differentiation factors (GDFs), Activins, Nodals and TGFβs. Upon ligand binding, a heterotetrameric receptor complex formation is initiated, composed of two type I and two type II serine-threonine kinase transmembrane receptors [20]. Type I receptors include the activin receptor-like kinases (ALK) 1-7, while type II receptors consist of the BMP receptor type 2 (BMPR2), Activin receptor type 2A (ACVR2A), Activin receptor type 2B (ACVR2B) and TGFβ receptor type 2 (TGFβR2) [21, 22]. Signal transduction proceeds via two major branches: Activins, TGFβs and Nodals primarily engage ALK4, ALK5 and ALK7, activating SMAD2/3; in contrast BMPs and GDFs bind ALK1, ALK2, ALK3 or ALK6 to initiate SMAD1/5 phosphorylation [23–26]. Ligand-bound type II receptors phosphorylate and activate type I receptors, which then recruit receptor-regulated SMADs via their L45 loop, a structural motif critical for SMAD isoform specificity [27]. Phosphorylated SMADs (pSMAD2/3 or pSMAD1/5) form heteromeric complexes with SMAD4 and translocate into the nucleus to regulate specific target gene expression downstream of the two branches [23–26].

The TGFβ family has been shown to play a pivotal role in placentation, however with conflicting results [4, 28]. Some studies suggest that TGFβ enhances EVT invasion and upregulates matrix metalloproteinases MMP-2 and MMP-9, while others describe inhibitory effects on trophoblast motility in response to TGFβ in cell line models [29–32]. Recent work has further refined this view, demonstrating that in human trophoblast stem cells (hTSCs) and first-trimester placentas, SMAD2 activation may inhibit, whereas SMAD3 may promote, EVT differentiation and invasion [33–35]. While the canonical TGFβ−SMAD2/3 has been extensively studied in this context, the contribution of SMAD1/5 signaling to trophoblast function remains poorly understood.

We have previously detected that TGFβ regulates the activation state of the endothelium via two opposing type I receptors and their associated SMAD cascades [36]. A dual receptor activation has also recently been described in the context of pulmonary arterial hypertension (PAH) [37] as well as in cancer cells, in which TGFβ-induced EMT requires combinatorial signaling via both SMAD3 and SMAD1/5. Interestingly, TGFβ-induced lateral signaling via transactivation of ALK5 to ALK2 led to a transient SMAD1/5 phosphorylation [38].

TGFβ1 as the most abundant and ubiquitously expressed isoform of TGFβ family ligands is widely expressed in the placenta and maternal tissue and has been implicated as a key factor in human trophoblast invasion [39], yet the molecular details of its signaling dynamics remain incompletely understood. In this study, we identified a previously unrecognized component of TGFβ signaling in trophoblast cells: in addition to SMAD2/3 phosphorylation, TGFβ1 induced a transient SMAD1/5 phosphorylation via the kinase activity of both ALK5 (*TGFBR1*) and ALK2 (*ACVR1*). This non-canonical pSMAD1/5 activation promotes expression of the EMT-associated transcription factor SNAIL and enhances trophoblast migratory capacity. Single-nucleus transcriptomics of healthy first trimester placentas revealed specific enrichment of *ACVR1*, *TGFBR1*, *SMAD1* and *SMAD5* at the transition zone from CTBs to EVTs and in turn *SNAI1* and *SNAI2* in EVTs, supporting a spatially restricted role of this signaling axis in early differentiation. Notably, both pSMAD1/5 and SNAIL expression were reduced in preeclamptic placentas compared to healthy donors, suggesting that disruption of this pathway may contribute to impaired EMT and shallow trophoblast invasion in placental disorders.

Together, these findings define a TGFβ1-induced ALK5/ALK2-SMAD1/5 signaling cascade as a critical regulator of trophoblast EMT with potential relevance to the pathogenesis of preeclampsia.

## RESULTS

### SMAD1/5 phosphorylation and SNAIL expression are reduced in preeclamptic placental samples

A defining feature of preeclampsia is the insufficient trophoblast invasion which could be attributed to a reduced EMT marker expression such as SNAIL and SLUG (**Fig. 1A**). We started out by addressing their expression levels in placental samples from healthy women (n = 5) given at the time of delivery and women suffering from mild (n = 5) or severe (n = 5) features of preeclampsia, according to the International Society for the Study of Hypertension in Pregnancy (ISSHP) guidelines, using immunohistochemical stainings (IHC). We focused on the part of the placental sample closest to the placental bed or where the placenta attaches to the uterus. Indeed, expression levels of SNAIL and SLUG were reduced in mild and even to a greater extent in severe preeclamptic placentas, indicating that the impaired trophoblast invasion is dependent on EMT (**Fig. 1B, Fig. EV1**).

TGFβ signaling has been shown to be a major inducer of EMT during embryonic development, cancer progression and fibrosis [19, 40–42]. Growing evidence highlights the importance of the TGFβ family in prenatal development and preeclampsia but published data on related expression patterns and functionalities are contradictory. Clarifying the molecular mechanisms underlying these signaling cascades is crucial to further understand healthy placentation and the pathogenesis of preeclampsia. We assessed the activation of the main branches of the TGFβ family i.e. pSMAD1/5 and pSMAD2/3 in placenta from healthy and preeclamptic pregnancies. Interestingly, in trophoblast cells, SMAD1/5 phosphorylation levels were reduced in samples from women with severe symptoms of preeclampsia while oppositely, SMAD2/3 phosphorylation levels were increased in samples from women with mild as well as severe preeclampsia symptoms (**Fig. 1C**).

### TGFβ1 induces both, a short-term SMAD1/5 and a long-term SMAD2/3 phosphorylation via different receptor complexes in trophoblast cells

Given the reduced SMAD1/5 phosphorylation and lower SNAIL expression observed in trophoblast cells in placental samples from preeclamptic women compared to healthy controls, we next investigated the molecular mechanisms regulating EMT in trophoblasts. As the TGFβ signaling cascade has been identified as the main inducer of EMT [12, 18], we elucidated the signaling pathway as potential inducer of EMT in trophoblasts. HTR8/SVneo cells were stimulated with TGFβ1 and analyzed by Western Blotting and immunofluorescent stainings (IF). Remarkably, TGFβ1 stimulation induced both SMAD2/3 and SMAD1/5 phosphorylation while BMP4 treatment resulted in SMAD1/5 activation only (**Fig. 2A**). This finding was further confirmed by the increased nuclear intensity of pSMAD1/5 upon TGFβ1 treatment (**Fig. 2B**). TGFβ1-induced pSMAD1/5 peaked after 45 min of stimulation, whereas pSMAD2/3 levels were more sustained over time (**Fig. 2C**). These results indicate that TGFβ1 induces both, a short-term SMAD1/5 and a long term SMAD2/3 phosphorylation in trophoblasts, which is in line with previous findings in cancer cells and endothelial cells [38, 43].

**Figure 2:**
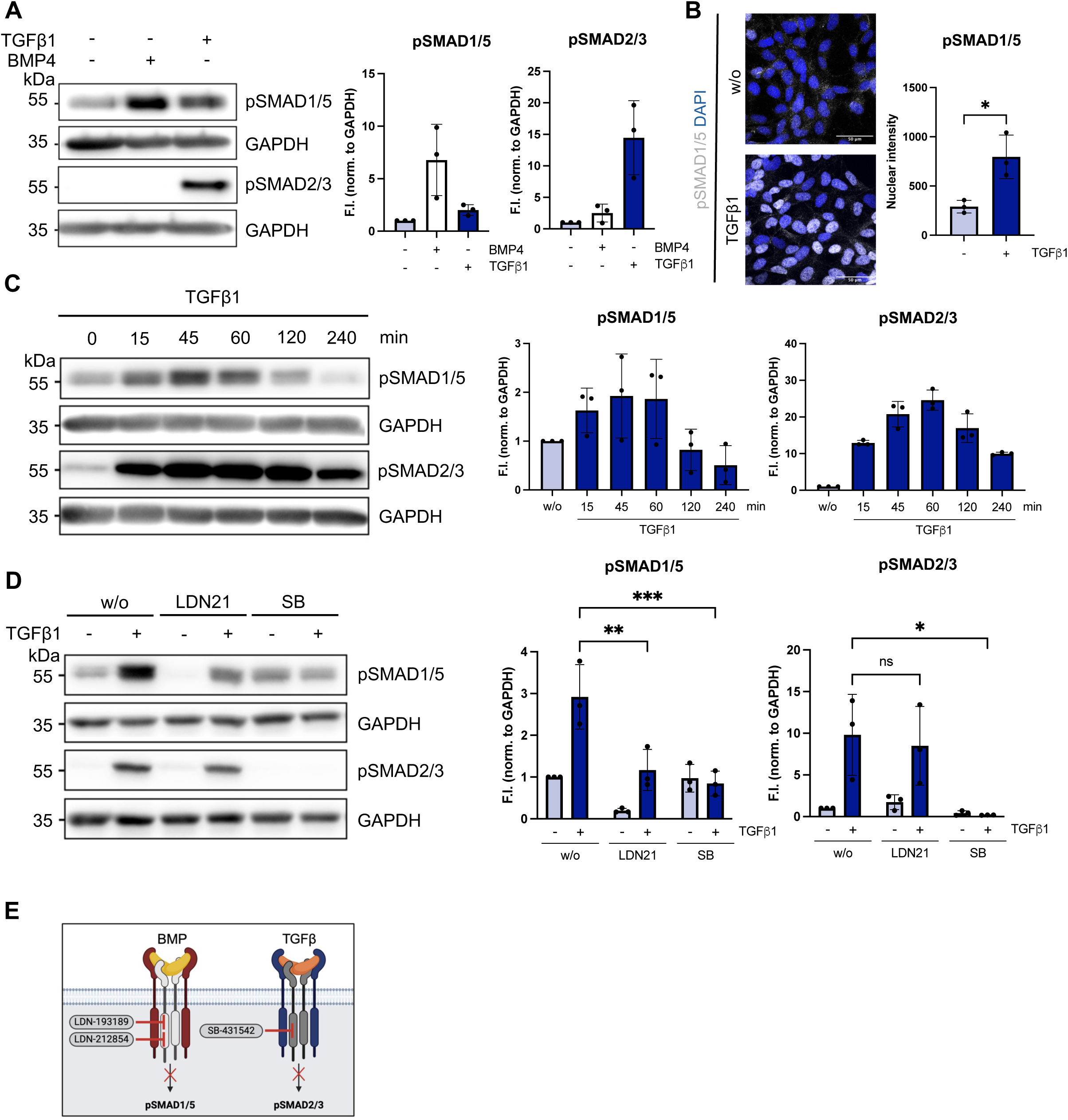
TGFβ1 induces both, a short-term SMAD1/5 and a long-term SMAD2/3 phosphorylation via different receptor complexes in HTR8/SVneo trophoblast cells. **A,C**, Representative immunoblots comparing HTR8/SVneo cells upon BMP4 (final conc. 25 ng/ml) or TGFβ1 (final conc. 10 ng/ml) stimulation for 45 min (**A**) or for indicated time points (**C**) depict the protein levels of pSMAD1/5, pSMAD2/3 and GAPDH as a loading control. **B**, Immunofluorescent stainings of HTR8/SVneo cells stimulated with TGFβ1 for 45 min show pSMAD1/5 (in grey) and DAPI (in blue). Nuclear intensity of pSMAD1/5 was calculated using ImageJ. Scale bar, 50 µm. **D**, Representative immunoblots comparing trophoblast cells upon inhibitor (LDN-212854 (final conc. 1 µM), SB-431542 (final conc. 10 µM)) and growth factor treatment depict the protein levels of pSMAD1/5, pSMAD2/3 and GAPDH. Quantification of protein expression is normalized to GAPDH and expressed as F.I. to w/o. Error bars represent the standard deviation from three independent experiments (n = 3). Statistical calculations relative to control are based on the unpaired t-test (**B**) and two-way ANOVA and Tukeýs post-hoc test, \**P* < 0.05, \*\**P* < 0.01, \*\*\**P* < 0.001 (**D**). **E**, Schematic overview of used inhibitors. ATP-competitive small molecule inhibitors target the kinase activity of the type I receptors. While the SB-431542 is a potent inhibitor of the TGFβ type I receptor, LDN-193189 and LDN-212854 have an inhibitory activity on BMP type I receptors.

Previous studies, including our own, have demonstrated that TGFβ can induce SMAD1/5 phosphorylation via distinct ALK receptors in various cancer, epithelial and endothelial cell types [38, 44–47]. In light of our observation that TGFβ1 stimulation leads to phosphorylation of both SMAD1/5 and SMAD2/3 in trophoblasts, we next sought to determine which receptor combinations mediate these signaling cascades. To address this experimental approach, different ATP-competitive small molecule inhibitors were used, targeting the kinase activity of the type I receptors and thereby preventing downstream SMAD phosphorylation (**Fig. 2E**). Interestingly, while treatment with LDN-193189 or LDN-212854 (BMP type I receptor inhibitors) led to reduced basal levels of pSMAD1/5, SB-431542 (TGFβ type I receptor inhibitor) completely abrogated pSMAD1/5 and pSMAD2/3 induction upon TGFβ1 stimulation (**Fig. 2D, Fig. EV2**). These results show that TGFβ1-induced SMAD2/3 and SMAD1/5 signaling are dependent on the kinase activity of a TGFβ type I receptor while the latter also indicates that a BMP type I receptor plays a contributory role.

### TGFβ1 signals through ALK2 and ALK5 to induce pSMAD1/5 and upregulate SNAIL in trophoblast cells

Since we observed a key function of the TGFβ type I receptor and a contributory role of the BMP type I receptor for TGFβ1-induced pSMAD1/5 signaling in trophoblasts, we next investigated the specific receptor combinations through which TGFβ1 exerts its effects. qPCR experiments revealed that *ACVR1* (encoding the BMP type I receptor ALK2) and *TGFBR1* (encoding the TGFβ type I receptor ALK5) are significantly expressed on RNA level in HTR8/SVneo cells (**Fig. EV3A**). Ramachandran et al. recently demonstrated a novel way of TGFβ-mediated receptor activation in cancer cells. TGFβ signaling occurs through the canonical TGFβ receptor ALK5 which in turn activates ALK2 upon ligand binding, leading to downstream SMAD1/5 phosphorylation which is required for EMT [38]. To further decipher the importance of the combinatorial kinase activity of two type I receptors in trophoblasts, we decided to focus on ALK2 and ALK5. Both receptors were knocked down individually in HTR8/SVneo cells, followed by TGFβ1 stimulation for 45 min to assess the transient SMAD1/5 phosphorylation. Notably, the pSMAD1/5 level was reduced upon siALK2 treatment while it was completly abolished in the absence of ALK5 compared to the control, suggesting that TGFβ1 signals initially through ALK5 to induce both pSMAD1/5 and pSMAD2/3 (**Fig. 3A, Fig. EV3C**). To further confirm the interplay between ALK5 and ALK2 upon TGFβ1 stimulation, HTR8/SVneo cells were adenovirally infected with dominant negative (dn) forms of both receptors. These kinase-dead mutants carry a lysine mutation (K233R), thereby preventing downstream SMAD phosphorylation. Overexpression of either dnALK2 or dnALK5, as well as their combined expression, resulted in a significant reduction of SMAD1/5 phosphorylation. In contrast, SMAD2/3 phosphorylation was significantly diminished only upon dnALK5 expression, indicating that the kinase activity of both receptors is required for TGFβ1-induced pSMAD1/5 signaling (**Fig. 3B**).

**Figure 3:**
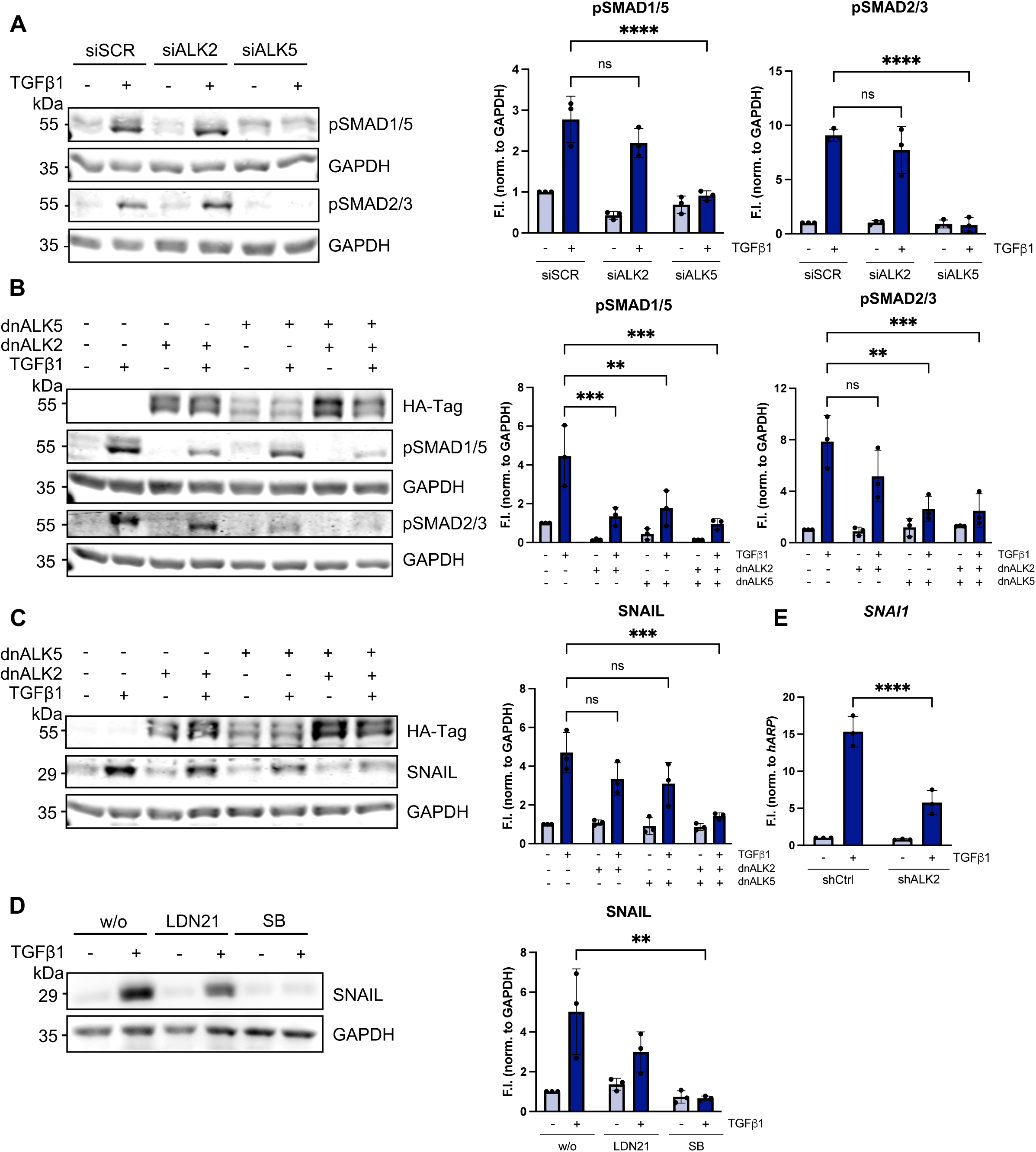
TGFβ-induced SMAD1/5 phosphorylation is dependent on an interplay between ALK2 and ALK5. **A**, Knockdown of ALK2 or ALK5 in HTR8/SVneo cells using siRNA. siSCR (scrambled) used as a negative control. Representative immunoblots comparing HTR8/SVneo cells upon TGFβ1 stimulation (final conc. 10 ng/ml) for 45 min depict the protein levels of pSMAD1/5, pSMAD2/3 and GAPDH. **B-C,** Adenoviral infection of HTR8/SVneo cells using LacZ (control), dnALK2 and dnALK5. Representative immunoblots comparing HTR8/SVneo cells upon TGFβ1 stimulation for 45 min (**B**) or 5 h (**C**) depict the protein levels of HA-tagged ALKs, pSMAD1/5, pSMAD2/3, SNAIL and GAPDH. **D**, Representative immunoblots comparing trophoblast cells upon inhibitor (LDN-212854 (final conc. 1 μM) or SB-431542 (final conc. 10 μM)) and growth factor treatment for 5 h depict the protein levels of SNAIL and GAPDH. **E**, Lentiviral infection of HTR8/SVneo cells using shControl and shALK2 constructs in pLKO.1 plasmids and puromycin selection (0.7 µg/ml). qPCR comparing expression of *SNAI1* in shControl and shALK2 cells upon TGFβ1 stimulation for 2 h. Quantification is normalized to GAPDH (**A-D**) or *hARP* (**E**) and expressed as fold induction (F.I.). Error bars represent the standard deviation from three independent experiments (n = 3). Statistical calculations relative to control are based on the two-way ANOVA and Tukeýs post-hoc test, \**P* < 0.05, \*\**P* < 0.01, \*\*\**P* < 0.001, \*\*\*\**P* < 0.0001.

Next, we examined whether the TGFβ1-induced interplay between ALK2 and ALK5 is required for downstream EMT marker expression in HTR8/SVneo cells, as previously shown in cancer cells [38]. Given that SNAIL acts primarily as a key inducer of EMT and its expression was reduced in preeclamptic placental samples (**Fig. 1B**), we focused subsequent analyses on this transcription factor. Remarkably, TGFβ1-induced SNAIL expression was reduced in trophoblasts overexpressing either dnALK2 or dnALK5 but significantly decreased when both dominant-negative receptors were co-expressed (**Fig. 3C**). Moreover, TGFβ1-induced SNAIL expression was decreased upon LDN-212854 treatment while it was completely abolished with the SB-431542 inhibitor (**Fig. 3D**). Furthermore, *SNAI1* expression was analyzed in stable ALK2 knockdown HTR8/SVneo cells using short hairpin ALK2 (shALK2). Upon TGFβ1 stimulation, *SNAI1* was significantly downregulated in the absence of ALK2 compared to shControl cells (**Fig. 3E, Fig. EV3D**). These data suggest that the TGFβ1-induced receptor interplay between ALK2 and ALK5 contributes to SNAIL upregulation which is required for trophoblast invasion.

### TGFβ1-induced pSMAD1/5 contributes to SNAIL expression

In addition to SNAIL (*SNAI1*), we studied the expression of other EMT transcription factors including SLUG (*SNAI2*) and TWIST (*TWIST*) as well as the mesenchymal marker N-Cadherin (*CDH2*) upon TGFβ1 stimulation in HTR8/SVneo cells. These factors were induced in a stepwise manner: *SNAI1* was upregulated as early as 2 h post-stimulation, followed by a peak in *SNAI2* expression and a concomitant increase in *CDH2* levels at 5 h. In contrast, *TWIST* expression did not show a significant change at the RNA level within 24 h (**Fig. EV4A**). SNAIL protein expression levels were significantly upregulated upon 5 h of TGFβ1 stimulation, as seen by Western Blotting and IF (**Fig. 4A,B**). Given that SNAIL is a known repressor of epithelial markers, including the transmembrane protein E-Cadherin (*CDH1*) [12, 18], we investigated the effect of TGFβ1 stimulation on *CDH1* transcriptional activity and E-Cadherin expression. As expected, HTR8/SVneo cells overexpressing the *hCDH1* promoter-tdTomato construct showed reduced *CDH1* transcriptional activity upon TGFβ1 treatment compared to control (**Fig. EV4C**). This result corresponded with decreased E-Cadherin expression, as confirmed by IF stainings (**Fig. EV4B**).

**Figure 4:**
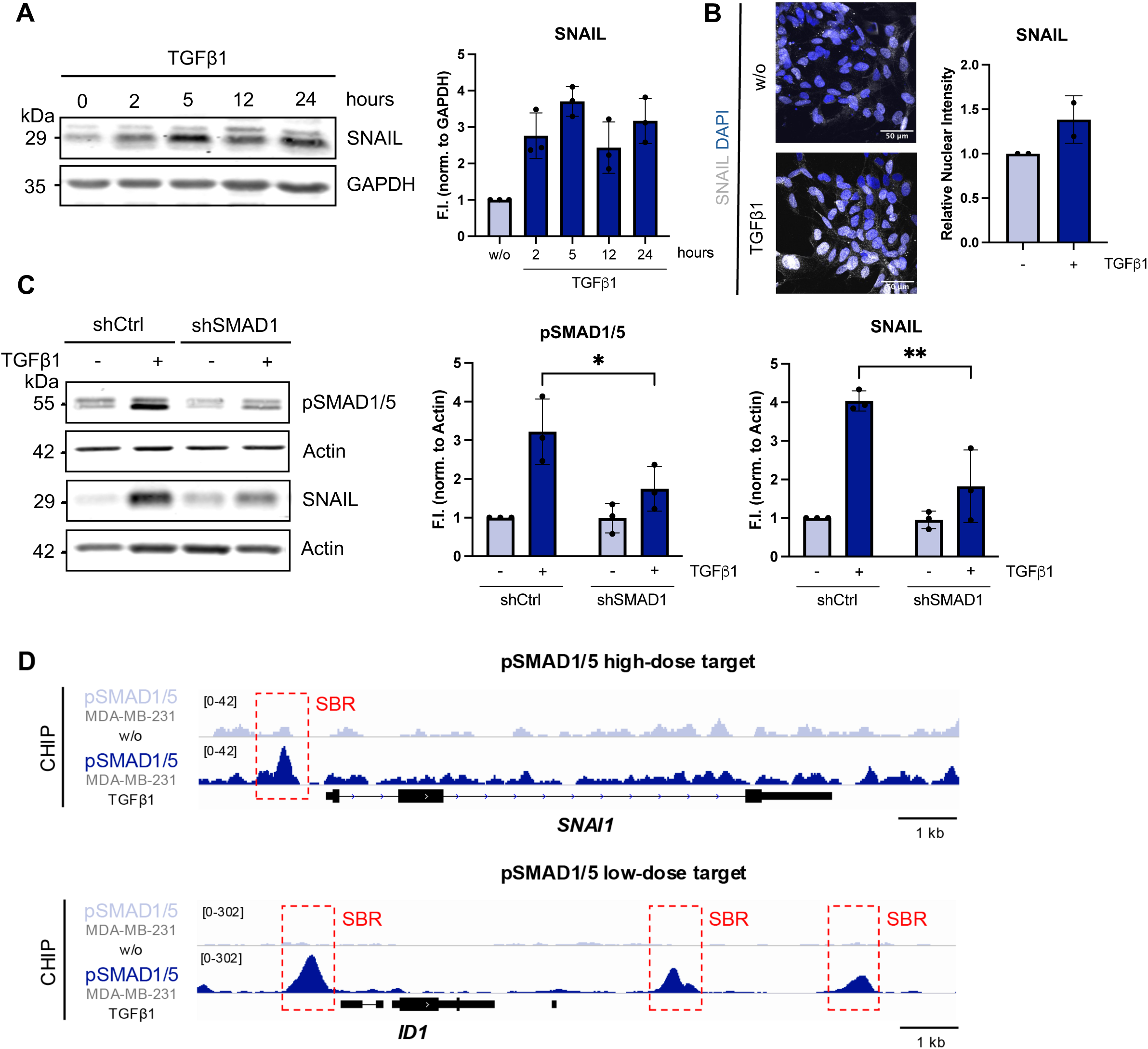
TGFβ-induced pSMAD1/5 contributes to SNAIL expression. **A,** Representative immunoblot of HTR8/SVneo cells upon TGFβ1 stimulation (final conc. 10 ng/ml) for the indicated time points depicts protein levels of SNAIL and GAPDH. **B**, Immunofluorescent stainings of HTR8/SVneo cells stimulated with TGFβ1 for 5 h show SNAIL (in grey) and DAPI (in blue). Nuclear intensity of SNAIL was calculated using ImageJ. Scale bar, 50 µm (n = 2 independent experiments). **C,** Lentiviral infection of HTR8/SVneo cells using shControl and shSMAD1 constructs in pLKO.1 plasmids and puromycin selection (0.7 µg/ml). Representative immunoblots comparing trophoblast cells upon TGFβ1 treatment for 45 min or 5 h depict the protein levels of pSMAD1/5 (45 min), SNAIL (5 h) and Actin. Quantification of protein expression is normalized to GAPDH (**A**) or Actin (**C**) and expressed as fold induction (F.I.) to w/o. Error bars represent the standard deviation from three independent experiments (n = 3). Statistical calculations relative to control are based on the two-way ANOVA and Tukeýs post-hoc test, \**P* < 0.05, \*\**P* < 0.01, \*\*\**P* < 0.001, \*\*\*\**P* < 0.0001. **D**, Genome browser view of publicly available TGFβ1-stimulated pSMAD1/5 ChIP-seq dataset (GSM2429820) on *SNAI1* and *ID1* loci. SBR – pSMAD1/5 bound region.

The link between the TGFβ1-pSMAD1/5 signaling pathway and the upregulation of the downstream EMT markers has not been shown before in trophoblasts. To further validate this, we generated stable HTR8/SVneo cells expressing short hairpin RNAs (shRNA) targeting SMAD1 (hSMAD1). In these cells, total SMAD1 protein levels were reduced by 50 % compared to control cells, confirming effective knockdown at the protein level (**Fig. EV4D**). Moreover, TGFβ1-induced pSMAD1/5 levels were reduced in shSMAD1 cells compared to shControl cells (**Fig. 4C**). Interestingly, the TGFβ1-induced expression of its downstream target SNAIL was significantly decreased in the absence of SMAD1, indicating that TGFβ1-induced pSMAD1/5 contributes to the upregulation of the EMT marker SNAIL (**Fig. 4C**).

To provide further insights into TGFβ1 regulation of SNAIL, we assessed publicly available ChIP-seq data of pSMAD1/5 in TGFβ1-stimulated MDA-MB-231 breast cancer cells (GSM2429820) [38] and identified TGFβ1-induced pSMAD1/5 bound regions in the promoter region of *SNAI1* and *ID1* compared to the untreated condition. Furthermore, *SNAI1* and *ID1* can be classified as pSMAD1/5 high-dose and low-dose targets, respectively, depending on the responsiveness to high or low pSMAD1/5 levels (**Fig. 4D**) [48].

Taken together, these findings indicate that TGFβ1-induced pSMAD1/5 transiently contributes to the upregulation of the EMT marker SNAIL.

### Trophoblast migration is promoted by TGFβ1 stimulation and ALK2

EMT has been correlated to increased cell migration and invasion, both characterized as hallmarks of the differentiation process in cancer cells [41, 49]. We used HTR8/SVneo cells with a stable knockdown of the EMT transcription factors SNAIL and SLUG to study their effect on trophoblast cell migration using a wound healing assay (scratch assay). Both *SNAI1* and *SNAI2* expression was decreased by 70 % upon TGFβ1 stimulation in the SNAIL and SLUG knockdown cells, respectively, compared to the non-silencing control cells (**Fig. EV5C,F**). The decreased EMT marker expression was accompanied by a reduced migratory capacity of trophoblasts, indicating that SNAIL and SLUG contribute to trophoblast migration (**Fig. EV5A-F**).

Since we have observed an upregulation of the EMT marker SNAIL in response to the TGFβ1-ALK2-ALK5-pSMAD1/5 signaling cascade, we next investigated its impact on trophoblast cell migration by performing wound healing assays with HTR8/SVneo cells cultured with or without TGFβ1 in the absence or presence of LDN-212854 or SB-431542. Notably, TGFβ1-stimulated cells migrated significantly faster compared to the untreated condition, while the migratory capacity was reduced upon either LDN-212854 or SB-431542 treatment (**Fig. 5A,B, Fig. EV5G**). To further confirm the importance of the BMP type I receptor ALK2 for trophoblast migration, HTR8/SVneo cells stably expressing shALK2 were used for the wound healing assay. Total ALK2 levels were reduced by 50 % in the respective shALK2 cells compared to shControl cells, confirming the knockdown efficiency on RNA level (**Fig. 5D**). In line with the LDN-212854-based results, the migratory capacity of trophoblast cells was significantly reduced in the absence of ALK2 compared to shControl cells (**Fig. 5C,D**).

**Figure 5:**
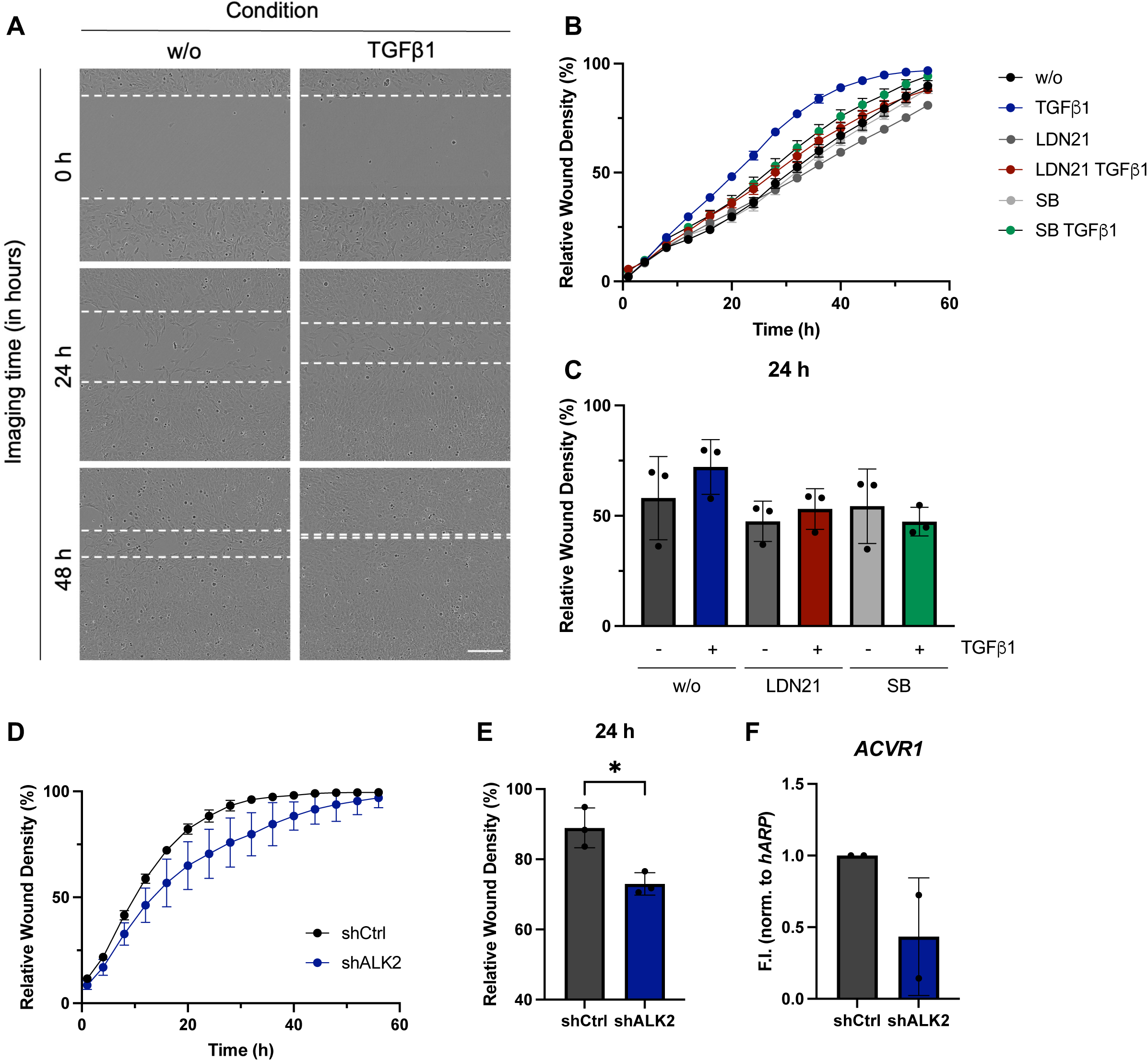
Trophoblast migration is promoted by TGFβ stimulation and ALK2. **A-C,** Migratory capacity of TGFβ1-treated HTR8/SVneo cells (final conc. 10 ng/ml) with or without inhibitors (LDN212854 final conc. 1 µM, SB431542 final conc. 10 µM) was assessed with the Wound Healing Assay. **A**, Images were taken over a time period of 56 h with 4 h intervals using the IncuCyte S3 Real-Time Quantitative Live Cell Analysis System. Dotted lines indicate the scratch. **B**, Wound closure (%) was plotted over time. Error bars represent the standard deviation from nine technical replicates (representative graph for three independent experiments). **C**, Wound closure (%) was plotted at the timepoint of 24 h. **D-F**, Lentiviral infection of HTR8/SVneo cells using shControl and shALK2 constructs in pLKO.1 plasmids and puromycin selection (0.7 µg/ml). **D**, Migratory capacity of succesfully transfected cells was assessed with the Wound Healing Assay. Cells were imaged for 56 h with 4 h intervals and wound closure (%) was plotted over time. Error bars represent the standard deviation from nine technical replicates (representative graph for three independent experiments). **E**, Wound closure (%) was plotted at the timepoint of 24 h. Statistical calculations relative to control are based on the unpaired t-test \**P* < 0.05. **F**, qPCR comparing expression of *ACVR1* in shControl and shALK2 cells. Values are normalized to *hARP* and expressed as fold induction (F.I.). Error bars represent the standard deviation from two (n = 2) (**F**) or three (**C,E**) independent experiments (n = 3).

To conclude, these findings demonstrate that the TGFβ1-induced pSMAD1/5 signaling cascade via a dual receptor activity of both ALK2 and ALK5 contributes to the upregulation of the EMT marker SNAIL that in turn facilitates trophoblast migration, a prerequisite for proper trophoblast invasion and conversion of the maternal spiral arteries.

### TGFβ1 signals through SMAD1/5 and SMAD2/3 phosphorylation to upregulate EMT markers in hTSCs

Trophoblast cell lines used to study human placental development, have been questioned with regard to their reliability and true relationship to the *in vivo* tissue state [50]. Recently, Okae et al. succeeded in deriving human Trophoblast Stem Cells (hTSCs) from the first-trimester placenta and the blastocyst [51]. The utility of these hTSCs for modelling trophoblast differentiation was confirmed by recapitulating functional phenotypes of primary trophoblasts and EVT subtypes and comparing them to corresponding first-trimester placental trophoblasts [52–54].

To evaluate if our novel findings can also be attributed to hTSCs, we used blastocyst-derived hTSCs as another cell model to study TGFβ-induced SMAD signaling and EMT. In line with previous experiments in HTR8/SVneo cells, TGFβ1 stimulation induced both SMAD2/3 and SMAD1/5 phosphorylation in hTSCs, while BMP4 treatment resulted in SMAD1/5 activation only (**Fig. 6A**). To examine the type I receptors required for SMAD signaling pathways, hTSCs were treated with different ATP-competitive small molecule inhibitors of type I receptor kinases. TGFβ1-induced pSMAD1/5 was significantly decreased upon LDN-212854 treatment while it was completly abolished when either SB-431542 or A83-01 (ALK4, 5, 7 inhibitor) were applied. In comparison, the pSMAD2/3 level was abolished only upon SB-431542 or A83-01 inhibitor treatment (**Fig. 6B**). To further decipher the importance of the combinatorial kinase activity of ALK2 and ALK5, as previously shown in HTR8/SVneo cells, hTSCs were adenovirally infected with either dnALK2 or dnALK5. Both type I receptors as well as their combined treatment led to decreased TGFβ1-induced SMAD1/5 phosphorylation while pSMAD2/3 levels were reduced only upon dnALK5 or the combined receptor overexpression (**Fig. 6C**). Thereafter, we studied the effect of TGFβ1 signaling on the EMT marker SNAIL and found that its expression was increased upon TGFβ1 stimulation whereas BMP4 had no effect on SNAIL expression (**Fig. 6E**). Furthermore, TGFβ1-induced SNAIL upregulation was decreased upon dnALK2 and dnALK5 as well as their combined treatment (**Fig. 6D**). These results strongly suggest that TGFβ1 signals through ALK5 and ALK2 to induce SMAD1/5 phosphorylation and to upregulate SNAIL in hTSCs which is consistent with our findings in HTR8/SVneo trophoblast cells.

**Figure 6:**
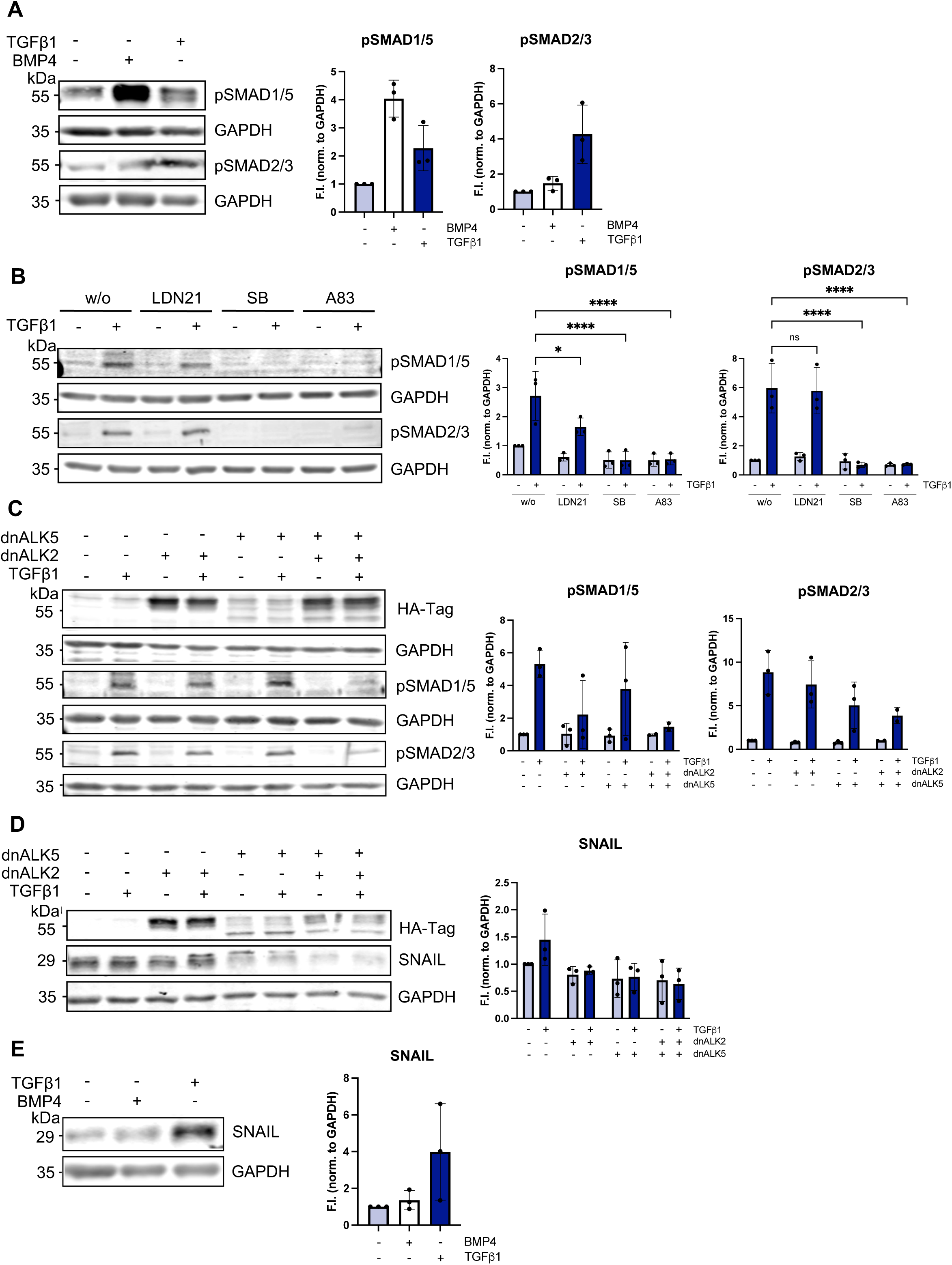
TGFβ signals through SMAD1/5 and SMAD2/3 phosphorylation to upregulate EMT markers in hTSCs. **A,E**, Representative immunoblots comparing human trophoblast stem cells (hTSC) upon BMP4 (final conc. 25 ng/ml) or TGFβ1 (final conc. 10 ng/ml) stimulation for 45 min (**A**) or 24 h (**E**) depict the protein levels of pSMAD1/5, pSMAD2/3 (**A**), SNAIL (**E**) and GAPDH (**A,E**). **B**, Representative immunoblots comparing hTSCs upon inhibitor (LDN-212854 (final conc. 1 μM) or SB-431542 (final conc. 10 μM) or A83-01 (final conc. 1 µM)) and growth factor treatment for 45 min depict the protein levels of pSMAD1/5, pSMAD2/3 and GAPDH. **C,D**, Adenoviral infection of hTSCs using LacZ (control), dnALK2 and dnALK5. Representative immunoblots comparing hTSCs upon TGFβ1 stimulation for 45 min (**C**) or 24 h (**D**) depict the protein levels of HA-tagged ALKs (**C,D**), pSMAD1/5, pSMAD2/3 (**C**), SNAIL (**D**) and GAPDH (**C,D**). Quantification of protein expression is normalized to GAPDH and expressed as fold induction (F.I.) to w/o. Error bars represent the standard deviation from three independent experiments (n = 3). Statistical calculations relative to control are based on the two-way ANOVA and Tukeýs post-hoc test, \**P* < 0.05, \*\**P* < 0.01, \*\*\**P* < 0.001.

### Single-nucleus transcriptomics reveal a specific ALK2/ALK5 and SMAD1/5 expression pattern at the transition area from distal CTBs to EVTs

To validate our findings in HTR8/SVneo cells and hTSCs and to investigate their clinical significance, we next analyzed the expression patterns of signaling components associated with the TGFβ-induced SMAD1/5 pathway in the publicly available single-nucleus RNA sequencing (snRNA-Seq) dataset. This dataset includes 10 first trimester placentas (5 – 11 weeks gestation) [55]. Nuclear identities within the dataset were grouped into clusters and visualized in a UMAP plot (**Fig. 7A**) as well as in a stream plot using the STREAM2 pipeline [56] (**Fig. 7B**). Distinct markers for specific trophoblast cell types were used to verify the stream plot of nuclear intensities in placental samples i.e. *TEAD4* (encoding TEA domain transcription factor 4) for proliferative CTBs, *ERVV-*1 (encoding Syncytin-2) and *ERVW-1* (encoding Syncytin-1) for pre-fusion CTBs and *HLA-G (encoding human leukocyte antigen G, HLA-G)* for EVTs (**Fig. EV6A**).

**Figure 7:**
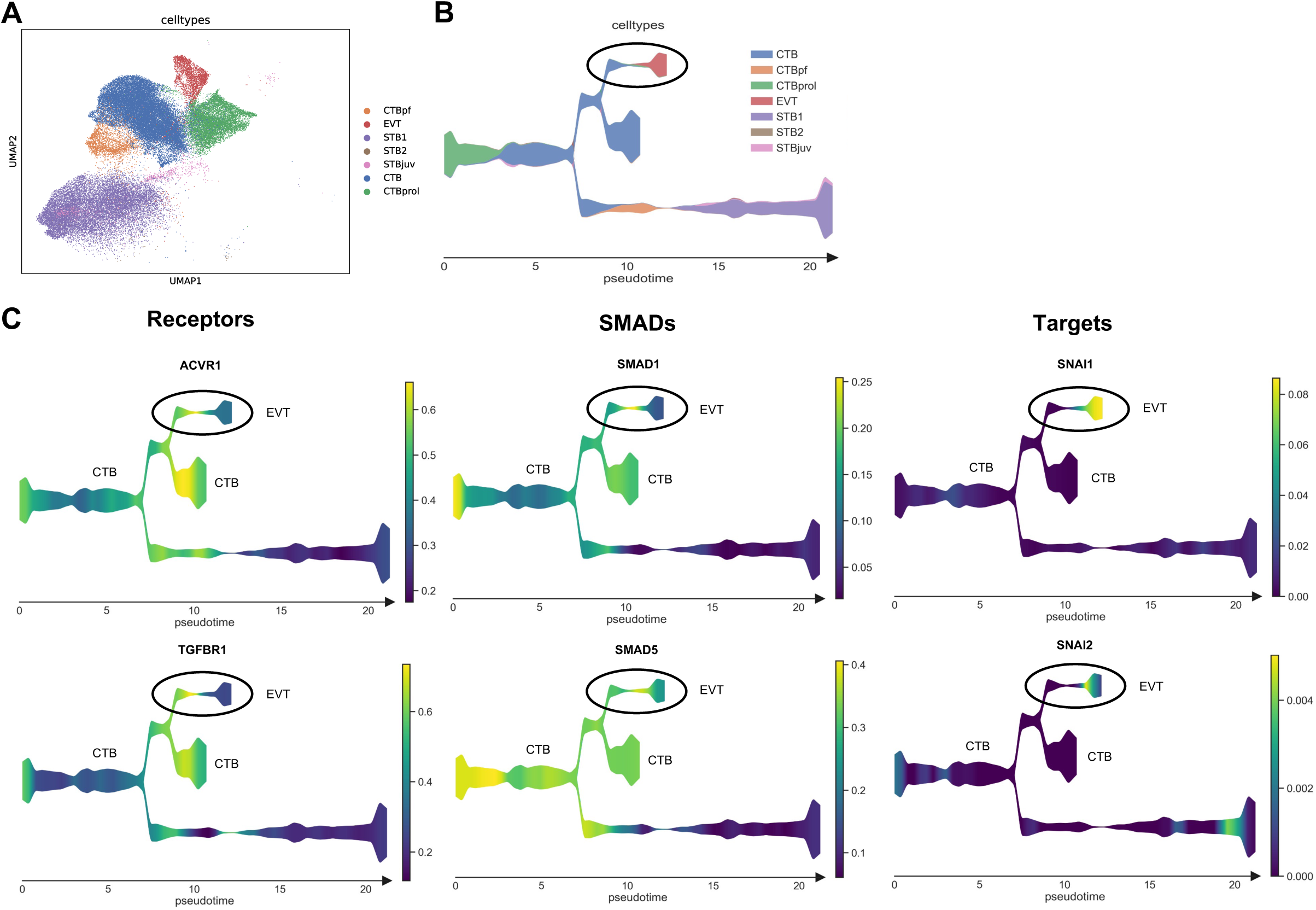
Single-nucleus transcriptomics reveals a specific ALK2/ALK5 and SMAD1/5 expression pattern at the transition area from CTBs to EVTs. Transcriptomic analysis of a publicly available snRNA-Sequencing dataset of first trimester placental tissues (n = 10, 5 – 11 weeks gestation) filtered for trophoblasts only (Nonn et al., *Hypertension* 2024). **A**, Analysis results in the following 7 nuclear identities visualized in a UMAP plot: proliferative cytotrophoblasts (CTBprol), cytotrophoblasts (CTB), pre-fusion cytotrophoblasts (CTBpf), juvenile syncytiotrophoblast (STBjuv), syncytiotrophoblast endocrine phenotype (STB1), syncytiotrophoblast mature phenotype (STB2) and extravillous trophoblasts (EVTs). **B**, Stream plot of nuclear identities in first trimester placental samples using the STREAM2 pipeline. CTBprol is the stream plot origin (time point = 0). The x-axis represents the pseudotime, reflecting the progression of cell differentiation, and the y-axis shows the gene expression levels along this trajectory. **C**, Stream plots of first trimester placental samples for expression of *ACVR1* (encoding ALK2), *TGFBR1* (encoding ALK5), *SMAD1*, *SMAD5*, *SNAI1* (encoding SNAIL), *SNAI2* (encoding SLUG).

Remarkably, both receptors *ACVR1* and *TGFBR1* as well as *SMAD1* and *SMAD5* were enriched at the transition area from CTBs to EVTs. While their expression decreased with the differentiation of EVTs, the EMT markers *SNAI1* and *SNAI2* increased and were specifically enriched in EVTs, along with the SMAD1/5 downstream target *ID1* (**Fig. 7C, Fig. EV6B**). *SMAD2* was widely expressed along the different trophoblast subtypes while *SMAD3* was significantly enriched in EVTs. However, both *SMAD2* and *SMAD3* had a low expression at the transition area from CTBs to EVTs compared to *SMAD1* and *SMAD5*. Furthermore, *SERPINE1* (encoding Plasminogen Activator Inhibitor-1, PAI-1), a downstream target of SMAD2/3 signaling, was not expressed in EVTs (**Fig. EV6B**).

Taken together, these observations strongly support that the TGFβ1/ALK2/ALK5/SMAD1/5 signaling pathway occurs transiently at the CTB/EVT interface and plays a key role in initial EVT differentiation by means of EMT.

## DISCUSSION

During the first trimester of pregnancy, fetal trophoblasts undergo EMT to penetrate the maternal decidua and remodel spiral arteries – an essential step for establishing high-to-low resistance for adequate placental perfusion. Dysregulation of this EMT process has been implicated in the pathogenesis of preeclampsia; however, the molecular mechanisms governing trophoblast differentiation at the fetal-maternal interface remain poorly understood. In the present study, we identify a previously unrecognized mechanism that enables human trophoblasts to undergo transient EMT during normal placental development. Using mutated forms of the ALK receptors along with selective small-molecule inhibitors targeting the ALK kinase function, we showed that transient TGFβ-induced phosphorylation of SMAD1/5 requires a cooperative signaling input from the canonical TGFβ type I receptor ALK5 and the BMP type I receptor ALK2. This dual-receptor signaling pathway promotes the expression of the EMT marker SNAIL and enhances trophobast migration, both critical for effective invasion of spiral arteries. Interestingly, snRNA-Sequencing data of first trimester placentas substantiates this model, revealing a spatially restricted enrichment of *TGFBR1*, *ACVR1*, *SMAD1* and *SMAD5* at the CTB to EVT transition zone. This expression pattern is consistent with localized, transient activation of the SMAD1/5 pathway during early stages of trophoblast invasion. Importantly, pSMAD1/5 and SNAIL expression levels were reduced in placental samples from preeclamptic pregnancies compared to healthy controls. Together, these findings support a model in which trophoblasts engage a finely tuned, transient TGFβ-induced SMAD1/5 signaling program to transiently adopt an EMT phenotype, which may subsequently facilitate mesenchymal-to-endothelial transition (MEndT) as cells progress towards a pseudo-endothelial identity, essential for spiral artery remodeling.

EMT has been extensively studied in the field of cancer progression and attempts have been made to further understand placental EMT by drawing parallels between both transition processes [15]. Based on common properties of cancer cells and trophoblasts, it is conceivable that both cell types undergo similar molecular mechanisms to induce distinct cellular responses such as EMT [16, 17]. Indeed, Ramachandran et al. described that TGFβ signals through both classes of type I receptors i.e. ALK2 and ALK5 to induce a transient SMAD1/5 phosphorylation in cancer cells, compared to a more sustained SMAD2/3 activation which occurs independently of BMP type I receptors. Using ChIP-Seq, the authors demonstrated that the initial transcriptional program is regulated by both SMAD signaling cascades to induce EMT and later on refined by the SMAD2/3 signaling pathway only [38].

The event of lateral signaling has not only been reported in different cancer cells but also in endothelial as well as distinct epithelial cells [38, 44–46, 57]. All these studies highlighted the importance of the ALK5 kinase domain activity for TGFβ-induced pSMAD1/5 signaling which is in line with our findings in trophoblast cells (**Fig. 2,3**). Goumans et al. demonstrated that in endothelial cells, ALK5 is required for the activation of ALK1, whereas in various cancer cell types, TGFβ stimulation enables ALK5 to activate ALK2 [38, 44–46, 57]. In addition to the required dual kinase activity, we observed a transient SMAD1/5 phosphorylation compared to a more sustained SMAD2/3 activation upon TGFβ stimulation which is consistent with previous studies in other cell models (**Fig. 2**) [38, 46, 47]. In contrast to our and previous publications, Liu et al. specifically showed that TGFβ-induced SMAD1/5 activation is critically dependent on the ALK5 L45 loop motif and occurs independently of BMP type I receptors, promoting mammary epithelial cell migration [58]. These findings appear contradictory to structural data indicating that ALK5 lacks an L45 loop motif compatible with the L3 loop of SMAD1/5, and to the well-established specificity of the ALK5 L45 loop for mediating selective phosphorylation of SMAD2/3. Nonetheless, the authors concluded that ALK5 plays a more direct function during TGFβ stimulation [58].

Several markers can be used to determine the occurrence of EMT, including morphological markers such as N-Cadherin, and EMT transcription factors like SNAIL and SLUG [59]. Those EMT markers were significantly increased upon TGFβ stimulation (**Fig. 4, Fig. EV4**), indicating that the TGFβ signaling cascade contributes to EMT in trophoblasts as it has been shown for cancer cells [12, 18, 38]. However, the EMT marker *TWIST* was not induced by TGFβ which is in line with findings by Okae et al. [51], assuming that the upregulation of *TWIST* might occur through a different signaling axis, such as the TNFα-induced NF-κB pathway [60]. The publications by Yu et al. and Tsubakihara et al. described that TGFβ signals through pSMAD2/3 to upregulate SNAIL in different cancer cells whereas Ramachandran et al. revealed that a combinatorial signaling via both pSMAD2/3 and pSMAD1/5 is required for TGFβ-induced EMT [12, 18, 38]. Supporting this dual-pathway model, Zeng et al. recently identified GDF15 as an upstream activator of the SMAD1/5 pathway, leading to increased EMT marker expression in HTR8/SVneo and JEG-3 trophoblast cells. Furthermore, the authors showed that *SNAI1* and *SNAI2* were transcriptionally regulated by SMAD5 which consequently promoted trophoblast cell migration and invasion [61]. In this study, we are showing for the first time that TGFβ-induced lateral signaling via ALK2 and ALK5 leads to transient SMAD1/5 phosphorylation and contributes to the expression of SNAIL, resulting in an increased migratory capacity of human trophoblast cells. However, high levels of pSMAD1/5 alone, as shown for BMP4, are insufficient to induce SNAIL, suggesting that both pSMAD1/5 and pSMAD2/3 are required for full transcriptional activation. High pSMAD2/3 and moderate pSMAD1/5 levels might form a certain threshold that has to be reached to fully upregulate SNAIL and thus induce EMT. Furthermore, pSMAD1/5 has been reported to form mixed heteromeric SMAD complexes [37, 62]. Concurrent activation of SMAD1/5 and SMAD2/3 may thus promote the formation of mixed SMAD complexes, such as SMAD3/3/1 replacing canonical SMAD3/3/4 complexes, potentially fine-tuning the signaling network upstream of SNAIL expression (**Fig. 8**).

**Figure 8:**
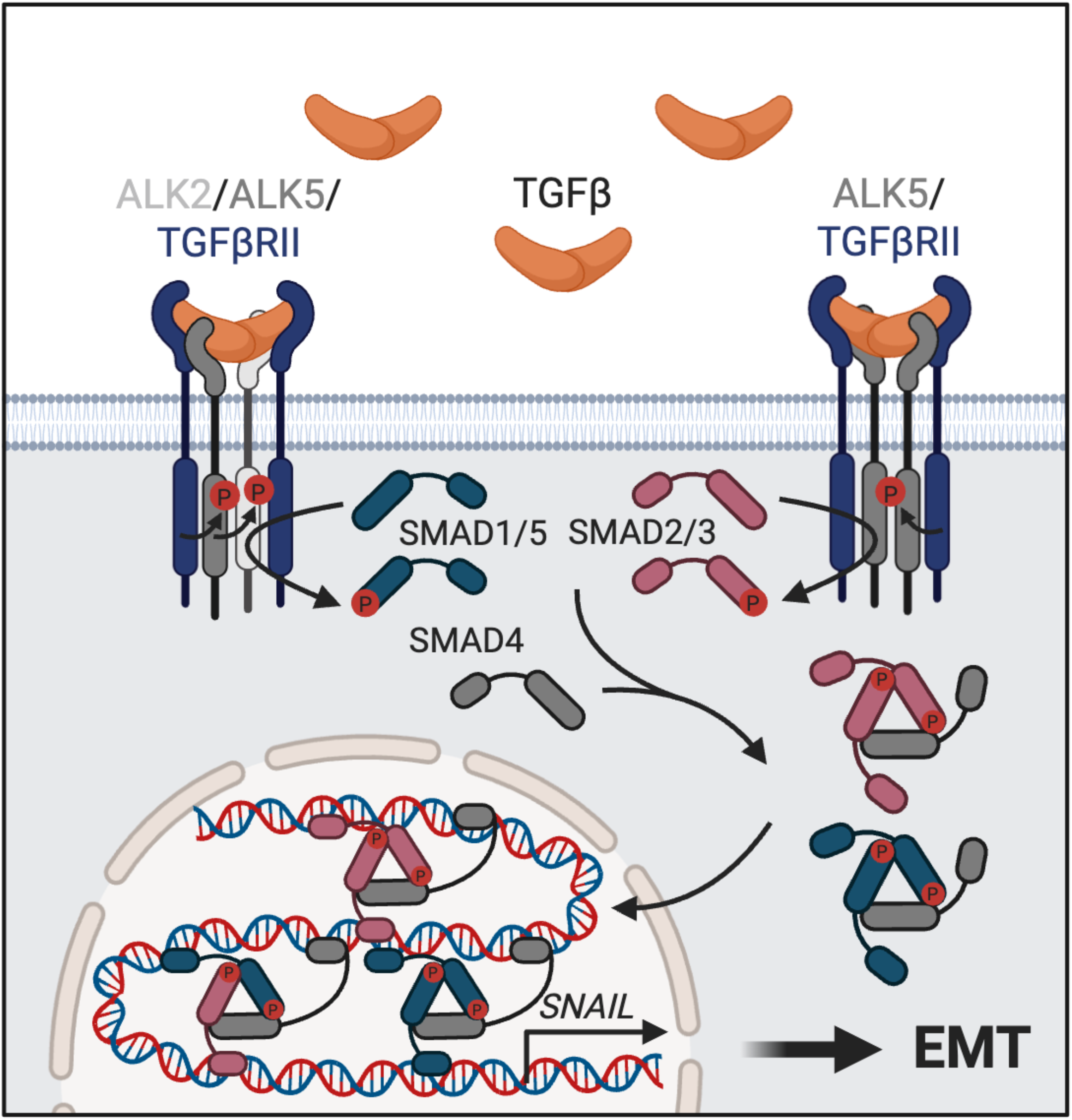
Overview of suggested signaling mechanism. TGFβ induces two distinct signaling cascades in trophoblast cells, a short-term SMAD1/5 phosphorylation via a kinase activity of both, the BMP type I receptor ALK2 and the TGFβ type I receptor ALK5, as well as a long-term SMAD2/3 activation independent of BMP type I receptors. TGFβ signaling through both SMAD pathways leads to a combined SMAD1/5 and SMAD2/3 regulation of downstream targets like the EMT marker SNAIL, a prerequisite to enter EMT.

The effect of TGFβ on cell migration and invasion seems to be contradictory throughout the literature. On the one hand, studies have described that TGFβ1 promotes trophoblast migration which is in line with our findings, while on the other hand several reports revealed an opposing effect on trophoblast migration as well as invasion (**Fig. 5**) [29–32, 63]. Notably, a recent study by Hashimoto et al. demonstrated that TGFβ1-induced SMAD2 signaling, which promotes the invasiveness of EVTs, is carefully regulated by the ERK pathway and SMAD1/5, and abnormalities in the interplay potentially contribute to excessive trophoblast invasion characteristic for the pregnancy disease placenta accreta [64]. Another study by Haider et al. further addressed a careful regulation of the TGFβ/SMAD2/3 signaling cascade, showing that it is not fully activated in the majority of placental EVTs to allow detachment and migration while upregulated TGFβ signaling leads to reduced migration and enhanced differentiation into interstitial EVTs in the decidua [34]. Taken together, these results highlight the importance of carefully regulated activation states of TGFβ-induced SMAD signaling pathways along the fetal-maternal interface, thus challenging experimental studies. Contradictory results might be related to different trophoblast cell lines which mimic certain phenotypes of STBs or EVTs, depending on the specific location of origin of the trophoblast cell lines used. Furthermore, different cell passage numbers and distinct TGFβ ligands as well as individual experimental approaches could cause conflicting experimental outcomes.

Collective cell migration is defined as the co-ordinated movement of multiple cells that still retain cell-cell contacts while undergoing collective polarization [65]. SNAIL has been shown to elicit collective migration in squamous cell carcinoma by upregulating the junctional protein claudin-11 [66]. Our results demonstrated that SNAIL was the first EMT transcription factor to be induced in trophoblast cells upon TGFβ treatment (**Fig. 4, Fig. EV4**). The upregulation of SNAIL via TGFβ-pSMAD1/5 signaling would perfectly fit the process of collective migration of the cytotrophoblast at the distal column of the anchoring villi.

To study human placental development and preeclampsia, different trophoblast models have been used, including primary trophoblasts, cell lines, placental explants, trophoblast stem cells, organoids as well as animal models [39]. In this study, the HTR8/SVneo cell line was used, derived from a first trimester placenta of a 6-to-12-week old embryo and transfected with simian virus 40 large T antigen (SV40T) [67]. Cell lines such as the HTR8/SVneo cell line are a good model to analyze dynamics as well as regulatory mechanisms such as TGFβ signal transduction in our study. Previously, Msheik et al. defined the HTR8/SVneo cell line as a model to investigate EMT during placentation. Interestingly, they highlighted the importance of the cell passaging number: higher passaging led to a more mesenchymal phenotype and different experimental outcomes, an observation that we also made with our cells [68]. Therefore, we used only HTR8/SVneo cells with low passaging numbers (p4 – p16) to conduct the experiments in this study. However, it has been shown that this cell line contains a heterogenous population of both epithelial-like trophoblasts and mesenchymal cells and it is controversial in different publications in how far they are truly representative of normal trophoblasts [69, 70]. To further evaluate our findings, we included blastocyst-derived hTSCs as another cell model to study TGFβ-induced EMT. hTSCs are more homogenous and have a similar gene expression profile as primary trophoblasts and allow cell fusion as well as differentiation, giving more insights in biological functions [39]. In line with our experiments in HTR8/SVneo cells, TGFβ-induced pSMAD1/5 signaling required a dual kinase activity of two type I receptors, i.e. ALK2 and ALK5 in hTSCs (**Fig. 6**).

The event of TGFβ-induced lateral signaling via ALK2 and ALK5 driving transient EMT in trophoblasts was supported by snRNA-Seq data of healthy first trimester placental tissues showing an enrichment of the signaling components at the transition area from CTBs to EVTs. While their expression decreased with EVT differentiation, *ID1* as well as *SNAI1* and *SNAI2* were specifically and gradually expressed (**Fig. 7, Fig. EV6**). Interestingly, pSMAD1/5 levels as well as SNAIL and SLUG were reduced in preeclamptic placental samples compared to healthy controls (**Fig. 1, Fig. EV1**). Zeng et al. revealed that SMAD5, SNAIL and SLUG were positively correlated with GDF15 and their expression levels were decreased in villous samples from women who experienced spontaneous abortion (miscarriage) compared to pregnant women who had a live birth [61]. In comparison to Zeng et al. we show for the first time reduced pSMAD1/5 as well as SNAIL and SLUG expression levels in preeclampsia, indicating that SMAD1/5 signaling is crucial during early pregnancy.

In contrast to *SMAD1/5*, *SMAD2* and *SMAD3* showed very distinct expression patterns along the trophoblast subtypes using snRNA-Seq data, which is in line with recent studies describing opposing effects of SMAD2 and SMAD3 on EVT differentiation and invasion (**Fig. EV6**) [33–35]. The authors revealed that SMAD3 was exclusively expressed in EVTs and regulated the expression of the EVT markers DAO and PAPP-A2 while SMAD2 exerted more inhibiting effects on EVT differentiation [33–35]. Notably, Brkic et al. demonstrated that pSMAD3 levels were significantly decreased in both early- and late-onset preeclamptic samples [33] whereas Xu et al. described a significant increase in pSMAD2 levels in early-onset preeclampsia, compared to healthy controls [71]. We observed elevated pSMAD2/3 levels in mild as well as severe preeclamptic samples, however we cannot distinguish between pSMAD2 and pSMAD3 due to the antibody used, which is the most commonly cited research antibody (**Fig. 1**).

Taken together, these findings signify that TGFβ-induced pSMAD1/5 signaling via ALK2 and ALK5 occurs transiently at the CTB/EVT interface to initially induce EVT differentiation whereas SMAD2/3 signaling has a longer-term effect that later refines the initial transcriptional program. Hence, we assume that a refined balance between SMAD signaling pathways upon TGFβ stimulation is critical to fully upregulate EMT markers required for successful EVT differentiation (**Fig. 8**). A shift in balance might lead to impaired trophoblast differentiation, contributing to different pregnancy diseases such as preeclampsia. Arutyunyan et al. has laid the foundation to resolve the differentiation trajectory of trophoblast cells [53], however, mechanisms as well as key proteins controlling the balance of SMAD signaling cascades at the fetal-maternal interface remain to be investigated to provide further insights into the network underlying healthy placentation.

## EXPERIMENTAL PROCEDURE

### Study subjects

Placental samples were collected from cases and controls at the Department of Obstetrics and Gynaecology at the National University Hospital of Iceland in collaboration with Professor Þóra Steingrímsdóttir and Associate-Professor Jóhanna Gunnarsdóttir. The sample collection was based on informed consent from all participants. Placental samples were given at the time of delivery with the approval of the Institutional Review Board of Landspítali – the National University Hospital of Iceland (Bioethics committee), permit (#48/2018) during 2020-2021. Participants were women diagnosed with preeclampsia (cases) and healthy controls. The diagnostic criterion for preeclampsia followed the International Society for the Study of Hypertension in Pregnancy (ISSHP) guidelines [72]. To be diagnosed with preeclampsia, a woman must have hypertension (³ 140 mmHg systolic blood pressure and/or ³ 90 mmHg diastolic blood pressure) and at least one of the following: proteinuria (ACR: Albumin creatinine ratio ³ 8 mg/mmol), fetal growth restriction (< 10^th^ percentile) or signs of organ dysfunction, such as observed neurological complications or abnormalities in the following blood tests: creatinine (> 90 µmol/L), thrombocyte count (< 150.000/µL), liver enzymes alanine aminotransferase and aspartate aminotransferase (> 40 IU/L). Cases were divided into subgroups with mild or severe features of the disease. Mild cases (n = 5) gave birth to normal weight infants at term (³ 37 weeks of gestation); however, significant proteinuria was present in all cases (median ACR 42.2; range 8.9 to 74.2 mg/mmol). Severe cases (n = 5) were all diagnosed preterm (range 31 + 6 to 36 + 3 gestational weeks + days). Median ACR was 72.6 mg/mmol, ranging between 38.5 to 490 mg/mmol. Four of the cases had severe hypertension (systolic blood pressure ³ 160mmHg) and the following severe features were observed: Fetal growth restricion (n = 2), abnormal liver function (n = 2), severe neurological symtoms (magnesium sulfate to prevent eclampsia, n = 2) and low platelets (n = 2).

The control group (n = 5) consisted of pregnancies without preeclampsia and classified as low-risk according to guidelines from the National Institute for health and Care Excellence (NICE) (https://www.nice.org.uk/guidance/ng133/chapter/Recommendations#reducing-the-risk-of-hypertensive-disorders-in-pregnancy); neither one high risk factor nor two moderate risk factor for preeclampsia. High risk factors: Previous history of gestational hypertension or preeclampsia, chronic hypertension, pre-gestational diabetes, kidney dysfunction or autoimmune disease. Moderate risk factors: first pregnancy, age > 40 years, > 10 years from last pregnancy, family history of preeclampsia, body mass index (BMI) > 35 kg/m^2^.

### Tissue collection

Placental samples were collected at the time of delivery following informed consent from all participants. Immediately post-delivery, biopsies measuring approximately 5 × 5 cm were excised from the placenta and fixed in formalin at 4°C for no longer than 1–2 days. Samples were then transferred to ethanol for storage prior to paraffin embedding. Each embedded tissue block was sectioned vertically into four slices, which were subsequently subdivided horizontally into four equal parts. For further analyses, the section closest to the placental bed was selected.

### Cell culture of HTR8/SVneo cells

Experiments were conducted using the human trophoblast HTR8/SVneo cell line from ATCC (CRL-3271) which were cultured in Gibco RPMI 1640 medium (11875093, Thermo Fisher Scientific) with 5% FBS (Thermo Fisher Scientific) and 1 % penicillin-streptomycin (full medium) at 37°C and 5 % CO_2_. For stimulation experiments, cells were transferred to a starvation medium 3h prior to the ligand stimulation, composed of Gibco RPMI 1640 medium with 0.5 % FBS and 1 % P/S (starvation medium).

### Cell culture of human trophoblast stem cells (hTSC)

hTSCs derived from the human blastocyst were a kind gift from Dr. Jacob Hanna. hTSCs were seeded on 1 % Matrigel and cultured according to the methods described by Oldak et al. [73]. For inhibitor experiments and adenoviral infections as well as for stimulations, hTSCs were cultured in hTSC full medium without the A83-01 inhibitor for 2 – 3 days prior to the experiments.

### Inhibitor treatment

For inhibitor experiments, LDN-193189 (final conc. 10 µM, Tocris Bioscience), LDN-212854 (final conc. 1 µM, Selleckchem) and SB-431542 (final conc. 10 µM, Tocris Bioscience) were added to the starvation medium and incubated at 37°C for 1 h prior to stimulation treatments. For hTSCs, the inhibitors were added directly to the hTSC full medium without the A83-01 inhibitor (no starvation).

### siRNA transfections

siRNAs were purchased from Dharmacon (Accell non-targeting control siRNA, Accell human ACVR1 siRNA-SMARTpool, Accell human TGFBR1 siRNA-SMARTpool). HTR8/SVneo cells were transfected using 5 µl Lipofectamine RNAiMAX (Thermo Fisher Scientific) with 40 nM siRNA in 6-well plates according to manufactureŕs instructions. 48 h after transfection, cells were starved and stimulated as described before.

### Ligand treatment

Stimulation experiments were performed with BMP4 (final conc. 25 ng/ml, Peprotech) or TGFβ1 (final conc. 10 ng/ml, Peprotech) for 45 min to investigate SMAD phosphorylation and for the indicated time points to study the EMT marker expression.

### Quantitative real-time PCR (qPCR)

Total RNA was isolated using the FastGene RNA Basic Kit (FG-80050, Nippon Genetics) according to manufactureŕs instructions. 1 µg of total RNA was reversely transcribed using the GoScript Reverse Transcriptase (Promega) according to manufactureŕs instructions and the RT-qPCR was conducted using the Maxima SYBR green/ROX qPCR master mix (Thermo Scientific) with primers (10 µM) listed in **Table 1**. All reactions were performed in triplicates, run on the Biorad CFX and further analysed with the Biorad CFX Manager. Target gene expression was quantified relative to *hARP*, *RSP9* or *ACTB* as housekeeping genes, based on the application of the ΔΔCt method.

**Table 1:**
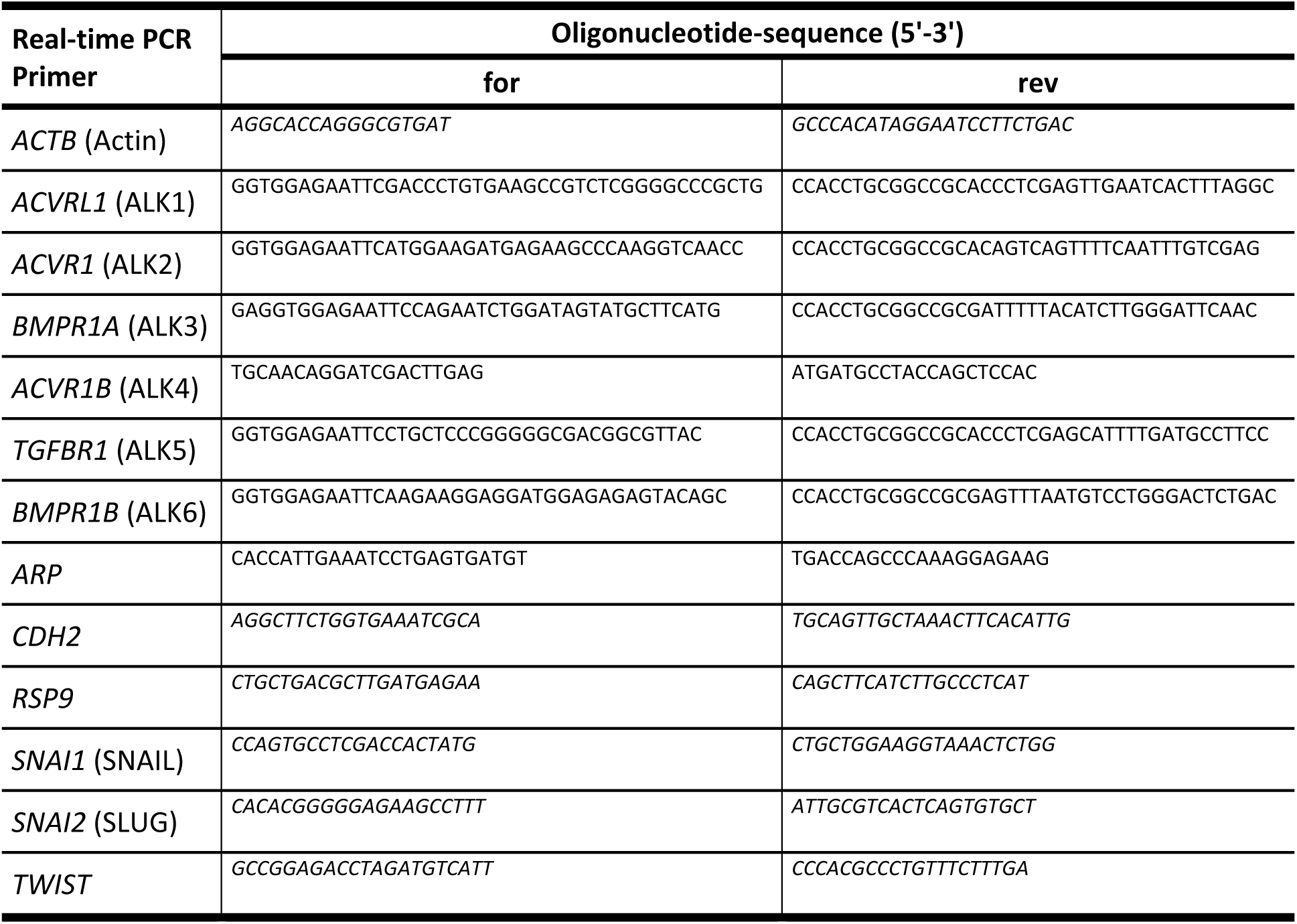
Oligonucleotides used for Quantitative Real Time PCR.

### Western Blot Analysis

Cells were lysed in Laemmli Buffer (1x) and boiled at 95°C for 7 minutes prior to loading on 10% SDS-PAGE gels. Proteins were transferred to a PVDF Immobilon-P membrane and the Western Blot was performed at 90 V for 2 ½ h. After blocking unspecific binding sites on the membrane with 3 % Bovine Serum Albumin (BSA) in Tris Buffered Saline with Tween-20 (TBS-T) for 1h at RT, membranes were incubated with primary antibodies at 4°C overnight. After TBS-T washing, membranes were incubated with secondary antibodies in blocking buffer for 1 h at RT, followed by further TBS-T washing and membrane detection using the Odyssey CLx Imaging System (LI-COR) (use of abcam pSMAD1/5, pSMAD2/3 from the Ludwig Lab and DyLight TM 680 secondary antibodies). For detection using the Fusion Imaging System, a 1:1 mixture of WesternBright Peroxide and WesternBright Quantum was added to the membrane to detect the enhanced chemiluminescence reaction (use of Cell Signaling pSMAD1/5 and pSMAD2/3 and the HRP secondary antibodies). Antibodies used for Western Blotting are listed in **Table 2** (GAPDH, HA-Tag and SNAIL antibodies were used for both detection techniques). The relative protein expression was normalized to GAPDH and analysed with ImageJ.

**Table 2:** Primary and secondary antibodies for Western Blotting.

| Antibody | Manufacturer | Species | Dilution | Type |
| --- | --- | --- | --- | --- |
| GAPDH | Cell Signaling (14C10) | Rabbit | 1:5000 | Primary |
| HA | Roche (12CA5) | Mouse | 1:1000 | Primary |
| SNAIL | Cell Signaling (C15D3) | Rabbit | 1:1000 | Primary |
| pSMAD1/5 | Abcam (ab76296) | Rabbit | 1:500 | Primary |
| pSMAD1/5 | Cell Signaling (41D10) | Rabbit | 1:1000 | Primary |
| pSMAD2/3 | Kind gift from Ludwig Lab | Rabbit | 1:1000 | Primary |
| pSMAD2/3 | Cell Signaling (138D4) | Rabbit | 1:1000 | Primary |
| mouse IgG, conjugated to HRP | Dianova/Jackson ImmunoResearch | Goat | 1:10.000 | Secondary |
| rabbit IgG, conjugated to HRP | Dianova/Jackson ImmunoResearch | Goat | 1:10.000 | Secondary |
| mouse IgG DyLight TM 680 conjugate | Cell Signaling (5366S) | Goat | 1:15.000 | Secondary |
| rabbit IgG DyLight TM 680 conjugate | Cell Signaling (5366S) | Goat | 1:15.000 | Secondary |

### Lentiviral Vector Production and Transduction of HTR8/SVneo

The pLKO.1 lentiviral cloning vector was used for cloning short hairpin RNA (shRNA) to generate lentiviral particles for the stable loss of function of ALK2 and SNAIL. HEK293T cells were transfected in a 6-well plate using FuGENE HD transfection reagent (Promega) and 10 µg DNA (3:1 ratio) according to manufacturer’s instructions. To generate lentiviral particles, a mixture of 400 µl of Opti-MEM Reduced Serum Medium (Gibco), 1,25 µg of pMD2 envelope plasmid (Addgene), 3,75 µg psPAX packaging plasmid (Addgene), 5 µg of the pLKO.1 shRNA plasmid (Sigma-Aldrich Mission shRNA human library (https://www.sigmaaldrich.com/catalog/search?term=mission+shrna+glycerol&interface=All&N=0&mode=partialmax&lang=en&region=IS&focus=product)) and 30 µl of FuGENE HD reagent was added to the cells and incubated at 37°C overnight. A control plasmid (p27) was used as shControl and a GFP containing plasmid (prRL) was used to check transfection efficiency. 3 days post-transfection, the lentiviral particles were collected, centrifuged and filter-sterilized through 0.45 µm pore-sized filters. HTR8/SVneo trophoblast cells were infected by adding 300 µl of the viral particles to the cells in full medium without P/S and 1,5 µl of polybrene for increased infection efficiency. 24 h after infection, the viral medium was replaced by fresh full medium and the cells were cultured further until a GFP signal was detectable. Subsequently, cells were treated with 0.7 µg/ml puromycin for the selection of cells stably expressing the puromycin-resistance cassette.

### Adenoviral infection

HTR8/SVneo cells were adenovirally infected with constructs expressing LacZ as a control and the dominant negative forms of ALK2 (dnALK2) and ALK5 (dnALK5) (a kind gift from Kohei Miyazono) in full medium. These kinase-dead mutants carry a lysine mutation (K233R), preventing downstream SMAD phosphorylation. 24 h after infection, the medium was replaced by fresh full medium and after further 24 h of culturing, starvation and stimulation was performed as described before.

### Transfection

HTR8/SVneo cells were seeded in 96-well Image Lock plates (Essen BioScience) and transfected at the same time with the *CDH1* promoter-tdTomato plasmid which was a kind gift from Dr. Aristidis Moustakas [74]. HTR8/SVneo cells were transfected using FuGENE HD transfection reagent (Promega) and 4 µg DNA (3:1 ratio) in Opti-MEM Reduced Serum Medium (Gibco) according to manufacturer’s instructions. The transfection mixture was incubated for 10 min at RT before adding it on top of the cells. 24 h after transfection, the cells were treated with small molecule inhibitors and growth factors, respectively (full medium +/- LDN-212854 (final conc. 1 µM), SB-431542 (final conc. 10 µM) and +/- TGFβ1 (final conc. 10 ng/ml)) and the total red object integrated intensity (µm^2^/image) was measured over a time period of 80 h with 2 h intervals using the IncuCyte S3 Real-Time Quantitative Live Cell Analysis System.

### *In Vitro* Migration assay

65.000 HTR8/SVneo cells were seeded in 96-well Image Lock plates (Essen BioScience) and cultured for 5 h. Cells were treated with inhibitors 1 h prior to the scratch assay as described before. Using the WoundMaker (Essen BioScience), a scratch was performed according to manufactureŕs instructions. Cells were washed three times with PBS and the respective cell medium was added (full medium +/- inhibitors and +/- TGFβ1 (final conc. 10 ng/ml)). Images were taken over a period of 48 h with 1 h intervals using the IncuCyte S3 Real-Time Quantitative Live Cell Analysis System as well as the corresponding software to detect and quantify the wound closure over time.

### Immunofluorescence Staining (IF)

For immunofluorescence stainings, cells seeded in gelatinized 12-well chamber slides (81201, Ibidi) were fixed with 2 % paraformaldehyde for 10 min and washed with PBS. Cells were permeabilized with 0.1 % Triton X-100, followed by blocking with 5 % goat serum and 1 % BSA in PBS (blocking buffer) for 1 h at RT. Subsequently, cells were incubated with primary antibodies (pSMAD1/5 abcam (ab76296) 1:250; SNAIL Santa Cruz (sc-271977) 1:250; E-Cadherin BD Biosciences (610181) 1:100) in blocking buffer in a humidity chamber at 4°C overnight. After PBS washing on the next day, the samples were incubated with secondary antibodies (goat α-rabbit Alexa488 and goat α-mouse Alexa647/Invitrogen, 1:1000) in blocking buffer in a humidity chamber for 1 h in the dark, followed by PBS washing and mounting using Fluoromount G with DAPI (00-4959-52, Invitrogen). Images were acquired with the FL 1200 Laser Scanning Confocal Microscope and the FV10-ASW Viewer Software (Ver.4.2b) and quantified using ImageJ.

### Image Analysis

The nuclear intensity of the pSMAD1/5 or SNAIL staining was calculated using the AnalyzeParticles Function in ImageJ. In short, the DAPI channel was selected and the background subtracted (*rolling = 200*). A threshold was created (*set Auto Threshold*) and converted to a mask, followed by the FillHoles function and AnalyzeParticles function (*size = 50 – Infinity display clear add*). After the background subtraction of the pSMAD1/5 or SNAIL channel, the nuclear ROIs from the DAPI channel were transferred to the pSMAD1/5 or SNAIL channel (*from ROI Manager*) and staining intensities were measured within the ROIs (*ROI Manager: Measure*).

### Immunohistochemical staining

Placental sections were immunohistochemically stained to assess both immunolocalization and expression of SNAIL and SLUG as well as phosphorylation of SMAD1/5 and SMAD2/3. The tissue samples were fixed in formalin, dehydrated in ethanol and embedded in paraffin blocks, sliced and placed on microscope slides. The tissue samples were deparaffinized in xylene and rehydrated in 90% ethanol, followed by boiling in Tris/EDTA-buffer pH 9.0 (10 mM Tris, 1 mM EDTA, 0.05% Tween-20) to expose the antigenic sites. Samples were blocked using peroxidase blocking buffer (ab64218, Abcam) for 5 min and further 10% FBS in PBS for 5-10 min at RT. Primary antibodies (listed in **Table 3**) diluted in 10% FBS/PBS, were added to the samples and incubated overnight at 4°C. After PBS washing the next day, samples were incubated with secondary antibodies for 30 min at RT. Subsequently, the DAB substrate chromogen solution (abcam (ab94665), 1:50 DAB chromogen in substrate solution) was added for 10 min. to form a brown precipitate, followed by hematoxylin staining for 10 sec for counterstaining. Coverslides were mounted using Eukitt quick-hardening mounting medium (Sigma-Aldrich, 03989) and imaged with the EVOS M7000 Imaging System.

**Table 3:** Primary and secondary antibodies for Immunohistochemistry.

| **Antibody** | **Manufacturer** | **Species** | **Dilution** | **Buffer** | **Type** |
| --- | --- | --- | --- | --- | --- |
| pSMAD1/5 | Abcam (ab76296) | Rabbit | 1:100 | TE | Primary |
| pSMAD2/3 | Invitrogen (44-244G) | Rabbit | 1:100 | TE | Primary |
| SNAIL | Santa Cruz (sc-271977) | Mouse | 1:100 | TE | Primary |
| SLUG | Santa Cruz (sc-166476) | Mouse | 1:100 | TE | Primary |
| Dako EnVision+ System – HRP labelled polymer, α-mouse/α-rabbit | Dako (K4001, K4003) | | | | Secondary |

### Single Nucleus RNA Sequencing (snRNA-Seq)

snRNA-Seq data was derived from a recent publication by Nonn et al. which describes the methods in detail [55]. 10 first trimester placental villi samples (5 – 11 weeks gestation) were included in this study and downstream analysis was performed as prior published. The dataset was filtered for trophoblasts only. Cluster annotations visualized in a UMAP showed the following nuclear identities: proliferative cytotrophoblasts (CTBprol), cytotrophoblasts (CTB), pre-fusion cytotrophoblasts (CTBpf), juvenile syncytiotrophoblast (STBjuv), syncytiotrophoblast endocrine phenotype (STB1), syncytiotrophoblast mature phenotype (STB2) and extravillous trophoblasts (EVTs). After nuclear cluster annotations, stream plots were generated for the target genes *ACVR1*, *TGFBR1*, *SMAD1*, *SMAD5*, *ID1*, *SNAI1*, *SNAI2*, *SMAD2*, *SMAD3*, *SERPINE1*, *TEAD4*, *ERVV1*, *ERVW1* and *HLA-G* using the STREAM2 pipeline (**S**ingle-cell **T**rajectories **R**econstruction, **E**xploration **A**nd **M**apping) where CTBprol was chosen as origin (time point = 0). The STREAM2 pipeline is available online (https://github.com/pinellolab/STREAM2) and the python package was used for the performance. The original STREAM pipeline was established and described by H. Chen et al. [56]. In these stream plots, the length of each branch represents the pseudotime, whereas the width is directly proportional to the number of cells at a given position. During the gene expression visualisation, the algorithm considers for each sliding window not only the number of cells but also their average gene expression values smoothed by bicubic interpolation (the maximum value is set as the ninetieth percentile of the average gene expression values from all the sliding windows). Detailed methods are described in the original publication [56].

### Statistical analysis

All experiments were performed in three independent biological experiments and the data were processed in GraphPad Prism version 10 for Apple (GraphPad Software, San Diego, CA, USA). The student’s t-test was used for comparing two groups with normally distributed data, while the one-way or two-way ANOVA was performed for multiple group comparisons. A significant difference was defined as p < 0.05.

## AUTHOR CONTRIBUTIONS

G.V. conceived the original idea and the project. S.M., H.I., M.A, and G.V. designed the study. S.M., H.I. and M.A performed the experiments and analyses. S.H. and O.N. provided the sequencing data and performed the sequencing analysis. Þ.S. and J.G. organized the collection of serum and placental samples at the University hospital. S.M. and G.V. wrote the original draft of the manuscript. J.J. and P.K. edited and critically revised the manuscript. All authors have read and agreed to the published version of the manuscript.

## ACKNOWLEDGMENTS

We are grateful to Dr. Kohei Miyazono for providing the adenoviruses and to Dr. P. ten Dijke for providing shRNA plasmids and to Dr. Aristidis Moustakas for the *CDH1* promoter-tdTomato construct. From Dr. E. Steingrimsson we appreciate helpful discussions. We thank Eydis Osk Simonardottir for assistance on *CDH1* promoter-tdTomato plasmid transfection and Bryndis Valdimarsdottir for expert technical assistance on immunohistochemical stainings. This research was supported by the Landspitali University Hospital Research Fund (grant no. A-2021-070) and a project grant (grant no. 228374-051) supported by the Icelandic Centre for Research (RANNIS). The German Research Foundation DFG supported this study by the grant SFB1444-TP04 to P.K.. The authors declare no potential/financial conflicts of interest related to this work.

## EXPANDED VIEW FIGURE LEGENDS

**Expanded View Figure 1 (Fig. EV1): SLUG expression is reduced in preeclamptic placental samples. A**, Immunohistochemical stainings of SLUG (counterstained with haematoxylin) on placental sections from healthy (Non-PE, n = 5) and preeclamptic women (PE) with mild features (n = 5). Immunostained sections were imaged with a light microscope using 10x and 40x magnification. Scale bar, 500 µm (10x) and 125 µm (40x). BV – blood vessel, EC – endothelial cells, TB – trophoblast cells, VS – villous space, IVS – intervillous space.

**Expanded View Figure 2 (Fig. EV2):** TGFβ1-induced SMAD1/5 phosphorylation requires the kinase activity of both BMP- and TGFβ type I receptors in HTR8/SVneo trophoblast cells. **A,** Representative immunoblots comparing trophoblast cells upon inhibitor (LDN-193189 (final conc. 1 µM), SB-431542 (final conc. 10 µM)) and growth factor (BMP4 (final conc. 25 ng/ml) or TGFβ1 (final conc. 10 ng/ml)) treatment depict the protein levels of pSMAD1/5 and Actin. Quantification of protein expression is normalized to Actin and expressed as F.I. to w/o. Error bars represent the standard deviation from three independent experiments (n = 3). Statistical calculations relative to control are based on the one-way ANOVA and Tukeýs post-hoc test, \*\*\**P* < 0.001, \*\*\*\**P* < 0.0001.

**Expanded View Figure 3 (Fig. EV3):** TGFβ-induced SMAD1/5 phosphorylation is dependent on an interplay between ALK2 and ALK5. **A,** qPCR comparing expression of BMP and TGFβ type I and type II receptors in HTR8/SVneo trophoblast cells. **B**, qPCR comparing expression of BMP and TGFβ type I receptors in HTR8/SVneo trophoblast cells to human umbilical arterial as well as human umbilical venous endothelial cells (HUAEC, HUVEC). **C**, Knockdown of ALK2 or ALK5 in HTR8/SVneo cells using siRNA. siSCR (scrambled) used as a negative control. qPCR comparing expression of *ACVR1* (encodes ALK2) and *TGFBR1* (encodes ALK5) shows a knockdown efficiency of 60 %. **D**, Lentiviral infection of HTR8/SVneo cells using shControl and shALK2 constructs in pLKO.1 plasmids and puromycin selection (0.7 µg/ml). qPCR comparing expression of *ACVR1* in shControl and shALK2 cells upon TGFβ1 stimulation (final conc. 10 ng/ml) for 2 h. Values are normalized to *RSP9* (**A,B**), *ACTB* (**C**) or *hARP* (**D**) and expressed as mean number of expression (MNE) (**A,B**) or fold induction (F.I.) (**C,D**). Error bars represent the standard deviation from three independent experiments (n = 3). Statistical calculations relative to control are based on the two-way ANOVA and Tukeýs post-hoc test, \**P* < 0.05, \*\**P* < 0.01, \*\*\**P* < 0.001, \*\*\*\**P* < 0.0001.

**Expanded View Figure 4 (Fig. EV4):** TGFβ signaling contributes to EMT marker expression. **A,** qPCR comparing expression of EMT markers *SNAI1, SNAI2, TWIST* and *CDH2* in HTR8/SVneo cells upon TGFβ1 (final conc. 10 ng/ml) stimulation for the indicated time points. **B**, Immunofluorescent stainings of HTR8/SVneo cells stimulated with TGFβ1 for 5 h show E-Cadherin (in grey) and DAPI (in blue). Staining intensity of E-Cadherin was calculated using ImageJ. Scale bar, 50 µm. Error bars represent the standard deviation from two independent experiments (n = 2). **C**, Transfection of HTR8/SVneo cells using the CDH1 promoter-tdTomato plasmid and treatment with TGFβ1 24 h post-transfection. Total red object integrated intensity (µm^2^/image) was measured over a time period of 80 h with 2 h intervals using the IncuCyte S3 Real-Time Quantitative Live Cell Analysis System and plotted over time (representative graph for n = 3 independent experiments) and at the time point of 60 h. **D**, Lentiviral infection of HTR8/SVneo cells using shControl and shSMAD1 constructs in pLKO.1 plasmids and puromycin selection (0.7 µg/ml). Representative immunoblots comparing trophoblast cells upon TGFβ1 treatment for 5 h depict the protein levels of SMAD1 and Actin as a loading control. Knockdown efficiency is represented by 50 % of shSMAD1 compared to shControl. Quantification is normalized to *ACTB* (**A**) or Actin (**D**) and expressed as fold induction (F.I.) to w/o. Error bars represent the standard deviation from three independent experiments (n = 3).

**Expanded View Figure 5 (Fig. EV5):** EMT markers SNAIL and SLUG are upregulated upon TGFβ1 stimulation and promote trophoblast migration. **A-F,** Lentiviral infection of HTR8/SVneo cells using shControl and shSNAIL (**A-C**) or shSLUG (**D-F**) constructs in pLKO.1 plasmids and puromycin selection (0.7 µg/ml). **A,D**, Migratory capacity of succesfully infected cells was assessed with the Wound Healing Assay. Cells were imaged for 56 h with 4 h intervals using the IncuCyte S3 Live Cell System and the wound closure (%) was plotted over time. Error bars represent the standard deviation from nine technical replicates (representative graph for three independent experiments). **B,E**, Wound closure (%) was plotted at the timepoint of 24 h. Error bars represent the standard deviation from three independent experiments (n = 3). **C,F**, qPCR comparing expression of *SNAI1* (**C**) and *SNAI2* (**F**) in shControl and shSNAIL or shSLUG cells upon TGFβ1 (final conc. 10 ng/ml) stimulation for 2 h. Values are normalized to *hARP* and expressed as fold induction (F.I.) (n = 1 independent experiments). **G,** Migratory capacity of TGFβ1-treated HTR8 cells (final conc. 10 ng/ml) with or without inhibitors (LDN212854 final conc. 1 µM, SB431542 final conc. 10 µM) was assessed with the Wound Healing Assay. Images were taken after 24 h and 48 h using the IncuCyte S3 Real-Time Quantitative Live Cell Analysis System and the dotted lines indicate the scratch.

**Expanded View Figure 6 (Fig. EV6):** Single-nucleus transcriptomics reveals distinct expression patterns of trophoblast markers, SMADs and the targets ID1 and SERPINE1. Transcriptomic analysis of a publicly available snRNA-Sequencing dataset of first trimester placental tissues (n = 10, 5 – 11 weeks gestation) filtered for trophoblasts only (Nonn et al., *Hypertension* 2024). Analysis results in the 7 following nuclear identities: proliferative cytotrophoblasts (CTBprol), cytotrophoblasts (CTB), pre-fusion cytotrophoblasts (CTBpf), juvenile syncytiotrophoblast (STBjuv), syncytiotrophoblast endocrine phenotype (STB1), syncytiotrophoblast mature phenotype (STB2) and extravillous trophoblasts (EVTs). **A,B**, Stream plots of nuclear identities in first trimester placental samples using the STREAM2 pipeline. CTBprol is the stream plot origin (time point = 0). The x-axis represents the pseudotime, reflecting the progression of cell differentiation, and the y-axis shows the gene expression levels along this trajectory. Stream plots of first trimester placental samples for expression of *TEAD4* (encoding TEAD4), *ERVV-1* (encoding Syncytin-2), *ERVW-1* (encoding Syncytin-1), *HLA-G* (encoding HLA-G) (**A**), *SMAD2*, *SMAD3*, *ID1*, *SERPINE1* (encoding PAI-1) (**B**).

